# Heteropentameric architecture predisposes functional inequivalence of acetylcholine receptor agonist sites

**DOI:** 10.1101/2025.09.13.676060

**Authors:** Johnathon R. Emlaw, Christian J.G. Tessier, Mariam Taktek, André R. Paquette, Christopher N. Boddy, Corrie J.B. daCosta

## Abstract

Muscle-type acetylcholine receptors are heteropentameric ion channels composed of four different but evolutionarily related subunits. These subunits assemble with a precise stoichiometry and arrangement, such that two distinct agonist-binding sites are formed at interfaces between a principal α-subunit and a complementary δ or ε/γ-subunit. Chemical differences between the two complementary subunits are presumed to confer functional inequivalence to the two agonist sites. This interpretation, however, overlooks the asymmetric architecture of the acetylcholine receptor, which places each subunit, and therefore each agonist site, at unique positions within the heteropentamer. The extent to which functional inequivalence of the agonist sites stems from this structural asymmetry, as opposed to chemical differences, remains unexplored. Here, we reconstruct an ancestral subunit capable of substituting for both complementary subunits, thereby engineering hybrid ancestral/human acetylcholine receptors with chemically identical agonist sites. Despite being chemically identical, these agonist sites remain functionally inequivalent. We show that this functional inequivalence stems from distinct intersubunit interactions dictated by the receptor’s heteropentameric architecture, which also underlie the subunit-dependent effects of a human disease-causing mutation. Thus, structural asymmetry emerges as a fundamental determinant of receptor function, capable of imposing functional inequivalence even in chemically identical sites, with implications for both the evolution of heteromeric protein complexes and the molecular basis of disease.

## Introduction

Nicotinic acetylcholine receptors are pentameric ligand-gated ion channels that mediate electrochemical signalling in the nervous system (Wonnacott et al., 2018). The nicotinic receptor superfamily has a rich evolutionary history, with vertebrate members being assembled from a diverse set of sixteen homologous subunits, related through a series of gene duplications (Le Novère and Changeux, 1995; Pedersen et al., 2019). The sixteen subunits are divided into α (α1-α7, α9, and α10), and non-α (β1-β4, δ, ε, and γ) subunits. While some α-subunits are capable of forming homopentamers (α7, α9, α10) (Couturier et al., 1990; Elgoyhen et al., 1994; Hone and McIntosh, 2022; Tekarli et al., 2025), most must assemble with non-α subunits within obligate heteropentamers as mixtures of two, three, or four different subunit types (Millar and Gotti, 2009; Gotti and Clementi, 2004; Zoli et al., 2015; Gharpure et al., 2020). The abundance of available subunits, and their ability to coassemble within heteropentamers, gives rise to a rich combinatorial repertoire of acetylcholine receptor assemblies, presumably evolved to meet a variety of synaptic needs (Sine and Engel, 2006).

Among vertebrate nicotinic receptors, the heteropentameric muscle-type acetylcholine receptor (AChR) exhibits the most subunit complexity, being the only member formed from four different subunit types. The adult form of the AChR assembles with two copies of the α1-subunit (α), and one each of the β1 (β), δ, and ε-subunits (Karlin et al., 1983). These five subunits come together in a counterclockwise α-ε-α-δ-β orientation, and are arranged like staves of a barrel around a central ion-conducting pore (Fig.1A) (Brisson and Unwin, 1985; Miyazawa et al., 2003). Both the stoichiometry and arrangement of these subunits is firmly entrenched, with the exception that the fetal γ-subunit replaces the ε-subunit during the early stages of development (Mishina et al., 1986; Missias et al., 1996). This subunit swap alters the unitary conductance of the channel, as well as its kinetic signature (Sakmann and Brenner, 1978; Mishina et al., 1986; Bouzat et al., 1994; Herlitze et al., 1996), directly demonstrating how subunit composition shapes AChR function, and thus the synaptic response.

**Figure 1.**
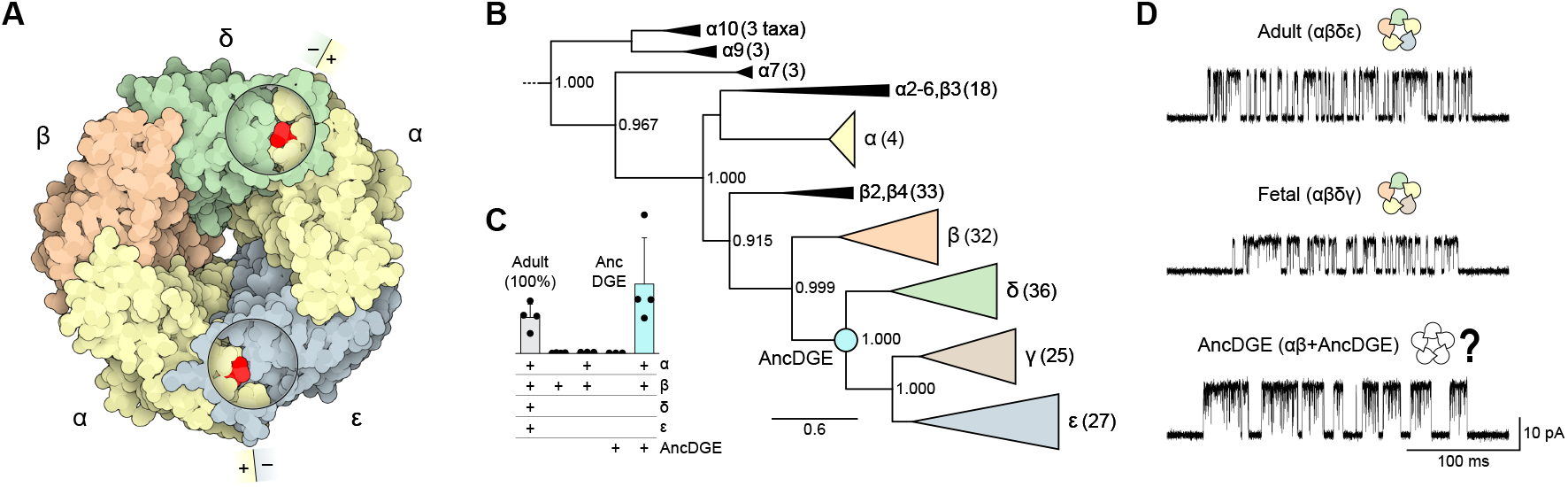
Reconstruction of AncDGE, a putative ancestor of the muscle-type acetylcholine receptor δ, γ, and ε-subunits. **(A)** Structure of the adult human AChR bound to acetylcholine (PDBID: 9DMH) (Li et al., 2025). Circular cutaways show the interfacial agonist sites, with acetylcholine in red. Plus (+) and minus (–) sides of each binding interface are indicated. **(B)** Cladogram of AChR paralogs informing the reconstruction of the muscle-type complementary subunit ancestor, AncDGE. The number of sequences for each paralog is included in parentheses. Internal labels are approximate likelihood ratios with the non-parametric Shimodaira-Hasegawa (SH) correction, and the scale bar represents the number of substitutions per site. **(C)** Binding of [^125^I]-α-Btx to cells expressing different combinations of AChR subunit cDNAs. Binding was normalized to cells transfected with the full complement of adult human AChR subunits (positive control), with cells transfected solely with the human β-subunit representing a negative control (n = 3-4). Mean plus standard deviation is shown alongside individual replicates. **(D)** Single-channel traces from cells expressing adult *(top)*, or fetal human AChRs *(middle)*, or cells transfected with human α, β, and AncDGE subunits *(bottom)*. Single-channel recordings were obtained in the presence of 30 µM acetylcholine at –120 mV, with openings shown as upward deflections. Traces filtered to 10 kHz. Subunits are colour-coded: α, yellow; β, orange; δ, green; γ, brown; ε, blue; AncDGE, cyan.

Acetylcholine activates the muscle-type AChR by binding to its two extracellular agonist-binding sites, which are located at the interfaces between one of the two α-subunits and either a δ or ε/γ-subunit (Blount and Merlie, 1989; Pedersen and Cohen, 1990; Sine and Claudio, 1991). Each agonist site is comprised of a series of loops, with residues from the α-subunits forming the principal (or plus/+) side, and residues from the non-α-subunits (δ, γ, or ε) forming the complementary (or minus/–) side. Divergence of the complementary δ, ε, and γ-subunits from a shared ancestor has rendered the two agonist sites chemically distinct (Fig.1A,B). It is reasonable to presume that these chemical differences also make the agonist sites functionally inequivalent. Indeed, unequal contributions of the two agonist sites have been reported on the basis of biochemical (Blount and Merlie, 1989; Pedersen and Cohen, 1990; Sine and Claudio, 1991; Kreienkamp et al., 1994; Papineni and Pedersen, 1997; Groebe et al., 1997; Prince and Sine, 1998b; Martinez et al., 2000; Molles et al., 2002a,b; Wang et al., 2003) and electrophysiological experiments (Sine et al., 1990; Zhang et al., 1995; Prince and Sine, 1998a; Sine et al., 2002; Tantama and Licht, 2009), the most convincing of which, showed that acetylcholine does not stabilize the open state equally when bound at either agonist site alone (Hatton et al., 2003; Shelley and Colquhoun, 2005; Shen et al., 2019). Attributing this functional inequivalence solely to the chemical differences of each agonist site overlooks an important consequence of the heteropentameric architecture of the AChR, which places each subunit in a unique environment within the pentamer. By virtue of their distinct interactions with different neighbouring subunits, each subunit position in the heteropentamer is structurally asymmetric. The extent to which the functional inequivalence of the two agonist sites can be explained by this structural asymmetry versus specific chemical differences is unknown. To address this we asked the deceptively simple question: what happens if we make the two agonist sites chemically identical?

Here we reconstruct a putative ancestor of the human δ, γ, and ε-subunits. We show that this ancestral subunit can substitute for both complementary subunits in hybrid ancestral/human AChRs, forming a ‘simpler’ receptor composed of three subunit types instead of four. As expected, this hybrid AChR possesses two chemically identical agonist sites, enabling a direct test of whether structural asymmetry alone can drive functional inequivalence. Mirroring the human AChR, single-channel recordings reveal that despite being chemically identical, the two agonist sites in hybrid ancestral/human AChRs are still functionally inequivalent. We further show that this functional inequivalence stems from distinct intersubunit interactions, which also explain the subunit-dependent effects of a human disease-causing mutation. These findings reveal that structural asymmetry within heteropentameric receptors can independently shape functional outcomes, even in the absence of chemical differences. This insight advances our understanding of how subunit arrangement, and specifically interactions at subunit interfaces, governs receptor function and susceptibility to disease.

## Results

We set out to engineer a muscle-type acetylcholine receptor with two copies of the same complementary subunit, instead of one each of the δ and ε (or γ) subunits. We reasoned that this ‘simpler’ AChR would have chemically identical agonist sites, making it possible to isolate how structural asymmetry arising from the heteropentameric architecture of the AChR contributes to the functional inequivalence of its two agonist sites. We began by inferring a molecular phylogeny of vertebrate acetylcholine receptor subunits, which delineates how the AChR subunits are related to one another, and at the same time represents a map for rationally exploring AChR subunit sequence space (Fig. 1B Fig. S1, S2 The phylogeny indicated that the three muscle-type complementary subunits (δ, ε, and γ) are more closely related to one another than they are to the other nicotinic receptor subunits. This suggests that their most recent common ancestor was a universal muscle-type complementary subunit, able to occupy both complementary subunit positions in an ancestral AChR. Based on this hypothesis we used ancestral sequence reconstruction (Hochberg and Thornton, 2017) to infer the maximum likelihood sequence of this putative ancestor, which we call ‘AncDGE’. AncDGE is a probabilistic inference based on [1] a multiple sequence alignment, [2] the presented molecular phylogeny, and [3] a best-fit model of amino acid evolution. AncDGE shares 62 % (Δ192 amino acid differences), 55 % (Δ225), and 51 % (Δ242) sequence identity with the human δ, γ, and ε-subunits, respectively (Fig. S3C). As a putative ancestor, AncDGE also exhibits sequence characteristics shared between, and unique to, the human δ, ε, and γ-subunits (Fig. S3D).

To assess whether AncDGE can substitute for the δ and ε (or γ) subunits we measured cell-surface binding of radiolabeled α-bungarotoxin (α-Btx) to cells transfected with combinations of cDNAs encoding the various AChR subunits (Fig. 1C). α-Btx is a competitive antagonist of muscle-type AChRs that binds to determinants on the α-subunit at the two agonist binding sites (Haggerty and Froehner, 1981; Gershoni et al., 1983; Rahman et al., 2020). Cells transfected with the full complement of adult human AChR subunits (α, β, δ, and ε-subunits) display robust α-Btx binding. Omitting the δ and ε-subunits eliminates α-Btx binding (Fig. 1C; Fig. S4). When AncDGE replaces both the δ and ε-subunits, robust α-Btx binding is restored, indicating that AncDGE can rescue cell-surface expression of human α and β-subunits. Single-channel recordings from cells transfected with α, β, and AncDGE cDNAs show that AncDGE-containing AChRs are functional, and qualitatively similar to those from cells expressing adult or fetal AChRs (Fig. 1D)

For AncDGE to replace both the δ and ε-subunits, there must be at least two copies of AncDGE in AncDGE-containing AChRs. To determine the subunit stoichiometry of AncDGE-containing AChRs we employed an electrical fingerprinting strategy (Emlaw et al., 2021 Electrical fingerprinting works by co-transfecting high-conductance (HC) and low-conductance (LC) variants of a particular subunit to yield channels whose single-channel amplitude depends on the number of HC versus LC variants incorporated. In effect, by tagging each AChR subunit with a reporter mutation that alters single-channel amplitude, we register the number of copies of each subunit type within the AChR pentamer. We began by fingerprinting the AncDGE subunit (Fig. 2A,B Table S1). We defined ‘wild-type’ AncDGE, which has a threonine at position 2^*′*^ (T2^*′*^) in the pore lining M2 helix, as our low-conductance subunit (AncDGE_LC_), and AncDGE with a glycine substituted at position 2^*′*^ (T2^*′*^G) as our high-conductance subunit (AncDGE_HC_). When AncDGE_LC_ cDNA is co-transfected with α and β-subunit cDNAs, the amplitude of channels expressed in each patch are uniformly distributed with a mean of 14.28 ± 0.41 pA (mean ± standard deviation; Fig. 2A,B middle). Co-transfecting AncDGE_HC_ with α and β-subunits also gives channels with a single, uniform amplitude, but with an increased mean of 17.64 ± 0.37 pA (Fig. 2A,B; top). When cDNAs encoding AncDGE_LC_ or AncDGE_HC_ are simultaneously co-transfected with α and β-subunit cDNAs a distribution of amplitudes are observed, all within the same patch (Fig. 2A; bottom). These amplitudes segregate into three distinct classes (Fig. 2B; bottom), with the highest and lowest amplitude classes mirroring the amplitudes observed when only AncDGE_HC_, or AncDGE_LC_, was included. The third class exhibits an amplitude intermediate to that of the HC and LC classes. Given that this intermediate class is only observed when both sets of cDNAs are present, the simplest interpretation is that channels with an intermediate amplitude have incorporated both types of subunits. Furthermore, because there is only one additional amplitude class, we conclude that channels with the intermediate amplitude class incorporate a single HC and a single LC subunit. By extension, low-amplitude channels contain two LC subunits, while high-amplitude channels contain two HC subunits, and thus AncDGE-containing AChRs contain two AncDGE subunits.

**Figure 2.**
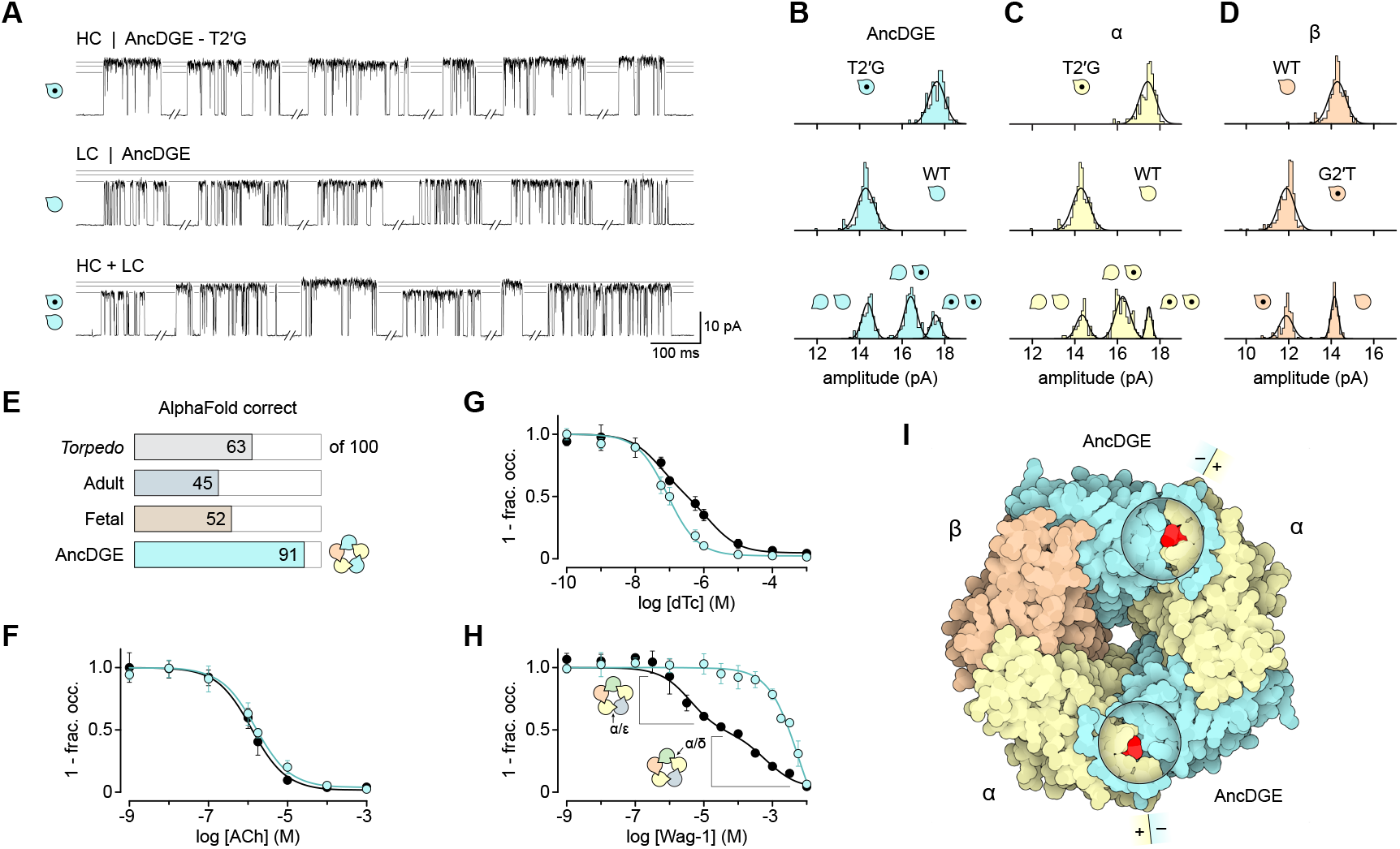
Subunit stoichiometry and arrangement of AncDGE-containing AChRs. **(A)** Representative single-channel bursts from cells co-transfected with cDNAs encoding wild-type human α, β, and either high-conductance (HC; *top*; T2*′*G), low-conductance (LC; *middle*; wild-type), or a mixture of HC and LC AncDGE subunits (HC + LC; *bottom*). Open subunits, and subunits with a dot, represent wild-type and conductance mutants, respectively. Bursts within each HC, LC, and HC + LC experiment originate from the same recording, and were elicited by 30 µM acetylcholine (at –120 mV). Bursts are filtered at 1 kHz for visual purposes (analyzed at 10 kHz). **(B-D)** Event-based amplitude histograms for HC, LC, and HC + LC experiments of AncDGE *(panel B)*, α *(panel C)*, and β subunits *(panel D)*. **(E)** AlphaFold3 predicted arrangement of *Torpedo*, human adult and fetal, and AncDGE-containing AChRs (see also Fig. S5). **(F-H)** Competition of [^125^I]-α-Btx binding with acetylcholine (ACh, *panel F* ), *d* -tubocurarine (dTc; *panel G*), and the Waglerin-1 peptide (Wag-1; *panel H*) to adult human (black) and AncDGE-containing AChRs (cyan). Data is shown as mean ± standard deviation from three transfections. **(I)** AlphaFold2 multimer model of AncDGE-containing AChRs. Circular cutaways show the interfacial agonist sites, with bound agonist in red.

We next extended the fingerprinting strategy to the α and β-subunits (Fig. 2C,D; Table S1). As with AncDGE, we identified HC and LC versions of each subunit whose amplitudes are uniformly distributed around a single mean value. Co-transfecting α_HC_ and α_LC_ cDNAs with those encoding the β and AncDGE subunits yields channels with three amplitude classes, mirroring that observed for AncDGE (Fig. 2C). Thus, following our interpretation above, we conclude that, like the human AChR, AncDGE-containing AChRs have two α-subunits. In contrast to AncDGE and α, when β_HC_ and β_LC_ are mixed, only two amplitude classes are observed (Fig. 2D). These two classes match the amplitudes from cells transfected with exclusively β_HC_ or β_LC_ (plus α and AncDGE cDNAs) indicating that only a single β-subunit is incorporated. Taken together, these data are consistent with a 2α:1β:2AncDGE subunit stoichiometry.

Electrical fingerprinting registers the number of copies of each subunit type, but does not tell us how these subunits are arranged within the heteropentamer. Given that AncDGE represents a putative ancestor of the δ, γ, and ε-subunits, a reasonable expectation is that the two AncDGE subunits replace the extant complementary sub-units in AncDGE-containing AChRs. We assessed whether the expected subunit arrangment of AncDGE-containing AChRs was plausible by predicting the subunit arrangement using AlphaFold3 (Fig. 2E; Fig. S5) (Abramson et al., 2024). The ability of AlphaFold3 to accurately predict subunit arrangements was first benchmarked using *Torpedo*, as well as both adult and fetal human AChRs, for which subunit stoichiometries and arrangements are well established. Known subunit stoichiometries were inputted and run 100 times for each set of sequences. In each case, only the top model from each run was considered (i.e., 100 total models for each set of sequences), with roughly half of all models registering the correct arrangement [63/100, *Torpedo*; 45/100, human adult; 52/100, human fetal] (Fig. 2E). Although a variety of alternate arrangements were also predicted (Fig. S5), the α and β-subunits were correctly placed in 70-80 % of models, indicating that the main struggle of AlphaFold3 was with correct placement of δ versus ε (or γ). We next predicted the subunit arrangement of AncDGE-containing AChRs based on the 2α:1β:2AncDGE stoichiometry determined in the fingerprinting experiments. In contrast to *Torpedo* and human AChRs, AlphaFold3 converged towards a single subunit arrangement in AncDGE-containing AChRs. Of the 100 models, 91 placed the two copies of AncDGE in the δ and ε locations, while also preserving α and β in their native positions (Fig. 2E; Fig. S5). Thus, AlphaFold3’s highest predicted arrangement for AncDGE-containing AChRs matches the expected counter-clockwise α-AncDGE-α-AncDGE-β orientation (Fig. 2E,I).

The highest predicted subunit arrangement by AlphaFold3 suggests that in AncDGE-containing AChRs the two agonist-binding sites are located at the two α(+)/AncDGE(–) subunit interfaces. If correct, the two agonist sites will each be composed of residues originating from the same two types of subunits, and should therefore be chemically identical. In the human AChR, several site-selective ligands display higher apparent affinity for one agonist site over the other, and thus expose chemical differences between them (Molles et al., 2002a; Wang et al., 2003). If the agonist sites in AncDGE-containing AChRs are indeed chemically identical, site-selective ligands should not be able to distinguish between them, and thus macroscopic binding of site-selective ligands is a useful way to assess agonist site chemistry.

To measure binding of various ligands, we took advantage of their ability to compete with radiolabeled α-Btx (Sine and Taylor, 1979). We began by measuring binding of acetylcholine to both adult human and AncDGE-containing AChRs (Fig. 2F; Table S2). Pre-incubating cells with increasing concentrations of acetylcholine inhibits α-Btx binding in a dose-dependent manner (Fig. 2F). For acetylcholine, binding is best-fit to a model with a single high-affinity acetylcholine binding site for both adult human and AncDGE-containing AChRs. Although acetylcholine does not distinguish between the two sites in these macroscopic binding experiments (Sine et al., 1994), this experiment does allow us to compare the relative apparent affinity of human and AncDGE-containing AChRs for acetylcholine. Evidently, despite 434 total amino acid differences between them, including several residues that map in and around the agonist sites (Fig. S3D), acetylcholine binding to the two types of receptors is essentially indistinguishable.

To evaluate whether the chemistry of the two agonist sites is the same, we turned to *d* -tubocurarine, whose site-selectivity is well-established (Blount and Merlie, 1989; Papineni and Pedersen, 1997). In adult human AChRs, *d* -tubocurarine binds with 15-fold higher affinity to the α/ε-site than to the α/δ-site (Wang et al., 2003). This preferential *d* -tubocurarine binding manifests as shallower binding in the competition experiment, which is best-fit with a two-site model (Fig. 2G; Table S2). By contrast, for AncDGE-containing AChRs, *d* -tubocurarine binding is as steep as acetylcholine binding, and best described by a single-site model (Table S2).

To further assess agonist site chemistry, we synthesized a ligand with a more pronounced site-selectivity. Waglerin-1 is a 22-amino acid peptide derived from the venom of *Tropidolaemus wagleri* (Wagler’s pit viper), which binds with 100 to 1000-fold higher affinity to the α/ε-site than to the α/δ-site (Molles et al., 2002b,a). The larger difference in affinity for the two agonist sites leads to a greater separation of their respective competition curves, leading to an almost stepwise decrease in binding, with each site contributing the expected 50 % of the total binding (Fig. 2H). As observed for both *d* -tubocurarine and acetylcholine, Waglerin-1 binding to AncDGE-containing AChRs is much steeper, and still best-fit to a model with a single type of binding site (Fig. 2H; Table S2). For both *d* -tubocurarine and Waglerin-1, fitting the binding data to a single-site model indicates that both ligands are unable to distinguish between the two agonist sites in AncDGE-containing AChRs, and thus they bind each site with similar apparent affinity. These data are consistent with AncDGE harboring chemically identical binding sites and corroborate the predicted counter-clockwise α-AncDGE-α-AncDGE-β arrangement, where ligands bind at two chemically identical α(+)/AncDGE(–) interfaces (Fig. 2I).

At the single-channel level, a hallmark of muscle-type AChR activity is that micromolar concentrations of acetylcholine elicit bursts of openings, occurring in quick succession and originating from the same channel (Sakmann et al., 1980). As the concentration of acetylcholine is increased, the probability of being open within these bursts also increases (Fig. 3A; Fig. S6, S7). For both human and AncDGE-containing AChRs, well-defined single-channel bursts can be measured at acetylcholine concentrations spanning 3 to 180 µM. Both AChRs exhibit a similar acetylcholine concentration dependence to their burst open probability (Fig. 3A). Kinetic analysis, through global fitting of the single-channel data to established schemes, reveals that despite differences in inferred rate constants, the equilibrium behaviour of both AChRs is quantitatively similar (Fig. S6, S7; Table S3). Specifically, the equilibrium constants describing acetylcholine binding or activation all differ by less than two-fold between human and AncDGE-containing AChRs (see ‘*K* _D1_‘, ‘*K* _D2_‘, and ‘Θ_1_‘, ‘Θ_2_‘ in Table S3). In particular, Θ_2_, which corresponds to activation of the di-liganded receptor is virtually indistinguishable between the two AChRs, thus it is not surprising that their single-channel dose responses resemble one another. Remarkably, despite 434 amino acid differences, distributed across the two complementary subunits, human and AncDGE-containing AChRs respond similarly to acetylcholine.

**Figure 3.**
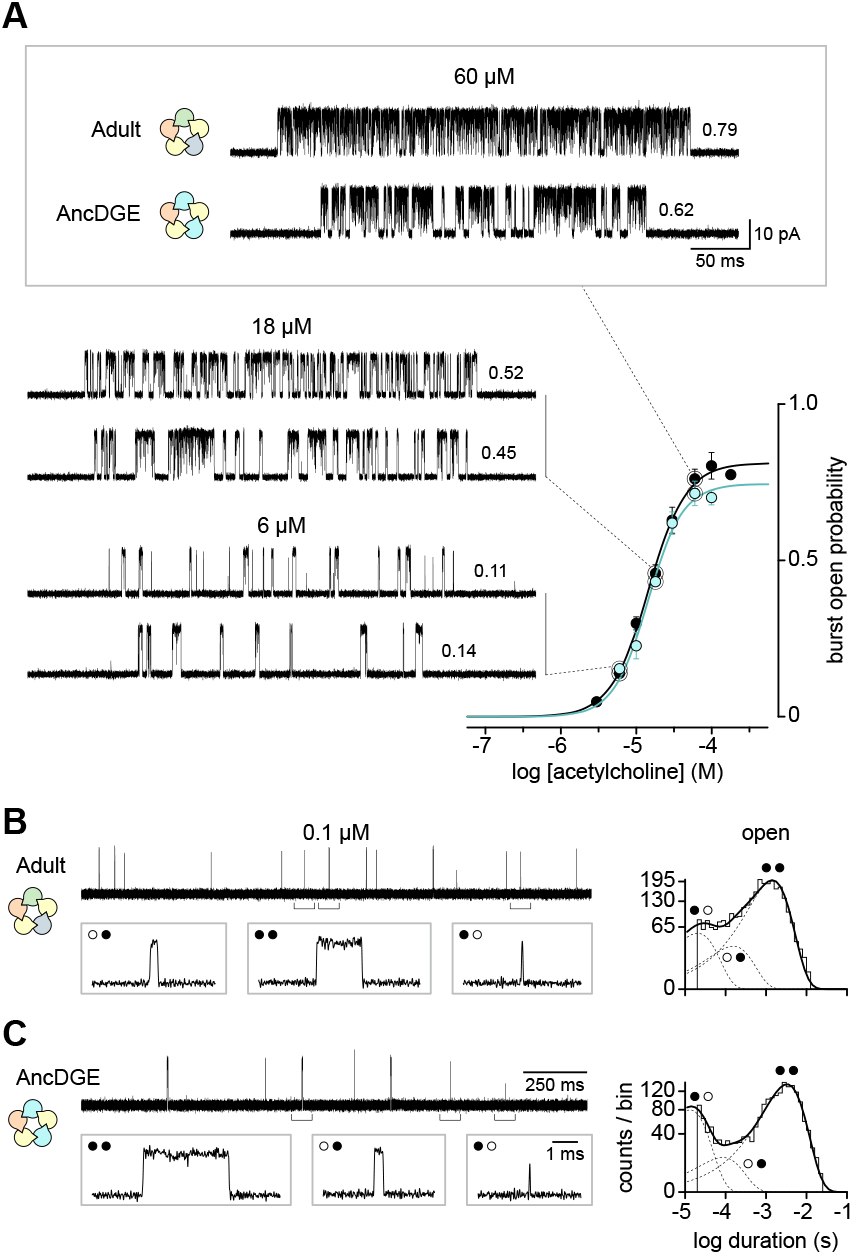
Single-channel activity of adult human and AncDGE-containing AChRs. **(A)** Dose-response of burst open probability as a function of acetylcholine concentration. Mean open probabilities are shown ± standard deviation (n = 3). Representative single-channel bursts (adult, *top*; AncDGE, *bottom*), and their corresponding burst open probability, are shown for the indicated acetylcholine concentrations (–120 mV; 10 kHz filter). **(B**,**C)** At low acetylcholine concentrations (0.1 µM), openings appear in isolation rather than in bursts. Recordings at 0.1 µM acetylcholine show three types of openings for adult human *(panel B)*, and AncDGE-containing AChRs *(panel C)*. Insets are an expansion of the bracketed regions, and show an example of each type of opening. Open-duration histograms were fit by a sum of three exponential components. Individual components are shown as dashed lines, and the summed fit as a solid line. Agonist occupancy levels for each class of opening are represented by circles (filled circle, occupied; open circle, unoccupied).

At acetylcholine concentrations below the threshold for eliciting bursts (i.e., <1 µM), single-channel openings occur infrequently, and in relative isolation (Fig. 3B,C). For the human AChR, the lifetimes of these isolated openings segregate into three components (Fig. 3B Fig. S8; Table S4). The longest-lived and most stable openings have been attributed to ‘di-liganded’ AChRs with acetylcholine simultaneously bound to both agonist sites, while the two classes of briefer openings are thought to originate from ‘mono-liganded’ AChRs with acetylcholine bound to either the α/δ or the α/ε-site (Hatton et al., 2003; Shelley and Colquhoun, 2005; Shen et al., 2019 The observation that there are two types of mono-liganded openings with distinct lifetimes indicates that agonists do not stabilize the open state equally when bound at either of the two sites. Thus, the two agonist sites are functionally inequivalent. Given that in the human AChR the two agonist sites are also chemically different, a reasonable hypothesis is that interactions between acetylcholine and agonist-site residues are different across the two sites, and thus the amount of energy available to stabilize the open state differs depending on whether the α/δ or the α/ε-site is occupied. This finding is consistent with cryo-EM structures, which show that agonists form distinct interactions in the two binding sites (Zarkadas et al., 2022; Li et al., 2024, 2025; Thompson et al., 2025).

If chemical differences between the two agonist sites account for the two distinct mono-liganded open state stabilities in the human AChR, then it seems reasonable to expect that AncDGE-containing AChRs, whose two agonist sites are chemically identical, should show a single type of mono-liganded openings. To our surprise, openings for AncDGE-containing AChRs still segregated into three exponential components (Fig. 3C; Fig. S8; Table S4). Mirroring human AChRs, AncDGE-containing AChRs displayed two types of brief mono-liganded openings, as well as a single class of longer di-liganded openings. Agonist occupancy levels were assigned on the basis that: [1] the two briefest components are absent at higher acetylcholine concentrations, which is a hallmark of mono-liganded AChR openings; and [2] the lifetime of the longest component matches that of openings occurring within bursts at higher acetylcholine concentrations. The distinct lifetimes of the two mono-liganded openings indicates that, despite being chemically identical, the two agonist sites in AncDGE-containing AChRs are still functionally inequivalent.

We hypothesized that the functional inequivalence of the two agonist sites is a consequence of AChR quaternary structure. Specifically, within the AncDGE-containing heteropentamer, the two AncDGE subunits are each sandwiched between different neighbouring subunits, which imparts them with their own unique set of intersubunit interactions. On their (–)-side, the two AncDGE subunits both interact with an α-subunit. On their (+)-side, the AncDGE in the δ-position interacts with the β-subunit, while the AncDGE in the ε-position interacts with another α-subunit (Fig. 4) We hypothesized that interactions across these distinguishing AncDGE(+)/β(–) and AncDGE(+)/α(–) interfaces could explain the functional inequivalence of the two chemically identical agonist sites.

**Figure 4.**
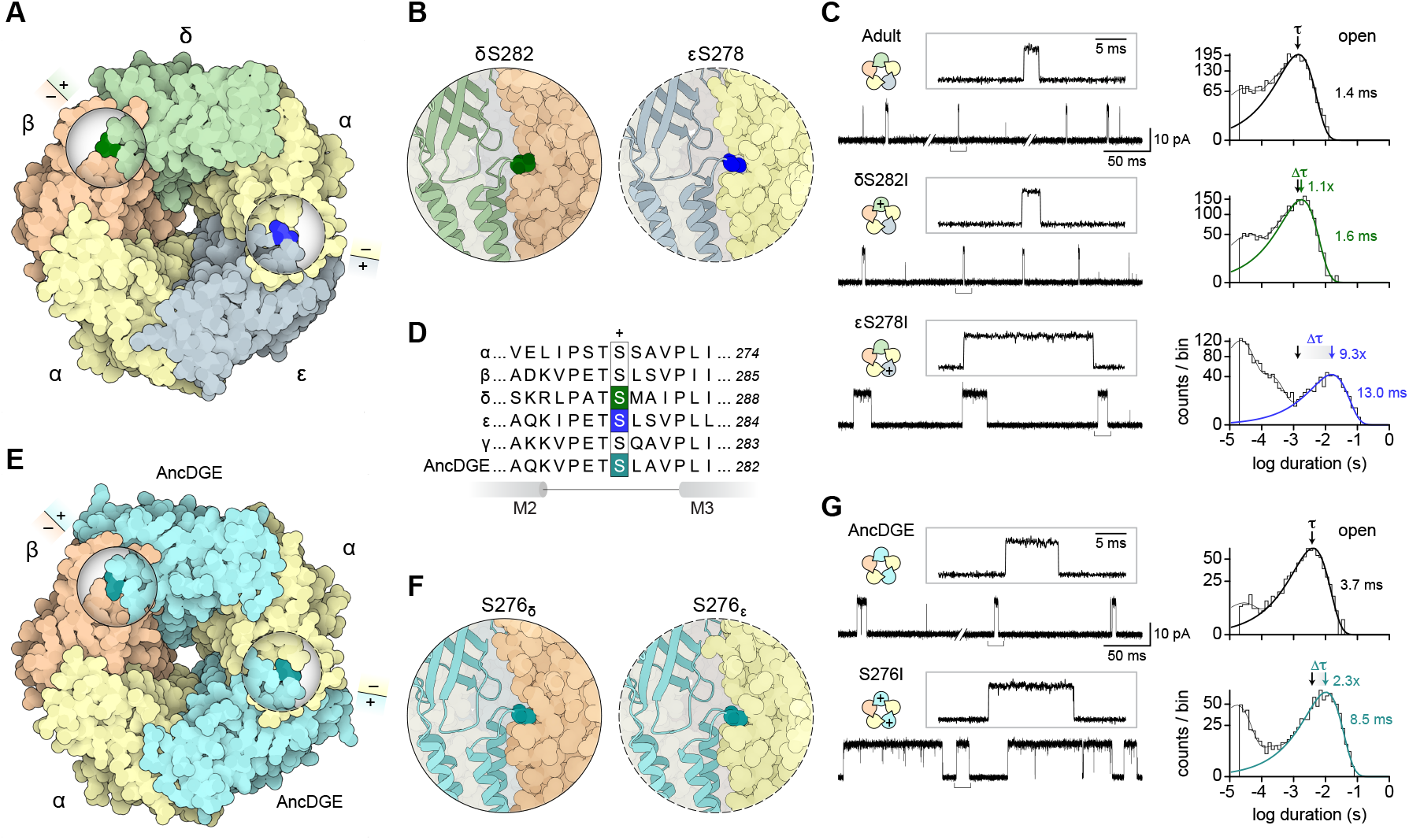
Subunit-dependent effects of a human disease-causing mutation. **(A)** Location of the serine-to-isoleucine substitution in δ and ε. Inset circles show cutaway views of the mutated site (bold colouring). (+) and (–) sides of subunit interfaces are indicated. **(B)** Close-up view of the mutated residue, as seen parallel to the membrane. The side chains of δS282 (*left* ; solid circle) and εS278 (*right* ; dashed circle) fit snugly against the (–) side of the β and α-subunit, respectively. **(C)** Single-channel recordings of wild-type and indicated serine-to-isoleucine substitutions. Plus sign (+) in each rosette denotes a mutant subunit. Individual openings (in the presence of 0.1 µM acetylcholine; –120 mV) are shown at two different time scales. Open-duration histograms are shown to the right, with the exponential component corresponding to the di-liganded lifetimes highlighted. Mean time constants for di-liganded fits, as well as the difference versus wild-type (Δτ), are indicated. **(D)** Multiple sequence alignment of the linker between M2 and M3 transmembrane segments, with the conserved serine that is mutated in each subunit highlighted. **(E**,**F)** Location of homologous substitutions in AncDGE-containing AChRs. **(G)** Single-channel activity of AncDGE and AncDGE-S276I. Openings were elicited by 1 µM acetylcholine at –120 mV.

To test this hypothesis, we exploited a previously characterized disease-causing mutation that maps to one of these distinguishing interfaces (Fig. 4A) Deletion of a single serine residue at position 278 in the human ε-subunit (εS278del) causes slow channel myasthenic syndrome by prolonging individual AChR openings (Fig. S9 Table S5) (Finlayson et al., 2013 A serine residue at this position is conserved in all human AChR subunits (Fig. 4D) and sub-stitution with isoleucine has been shown to similarly prolong openings of the mouse AChR, but in a subunit-dependent manner (Grosman et al., 2000). Of particular interest here, homologous substitutions in the δ and ε-subunits have divergent effects. Mutating the δ-subunit (δS282I) has negligible effect on open-channel lifetimes, while mutating the ε-subunit (εS278I) prolongs openings ∼9-fold, mimicking the effects of the original deletion/disease-causing mutation (Fig. 4A-C; Fig. S9; Table S5). While these divergent effects could stem from interactions at the distinguishing interfaces as hypothesized, it is also possible that amino acid differences between the δ and ε-subunits are responsible.

To eliminate the confounding differences in amino acid sequence between the δ and ε-subunits, we introduced the homologous substitution into AncDGE (AncDGE-S276I), and examined its effects in AncDGE-containing AChRs. Installing the S276I substitution into AncDGE leads to a 2.3-fold prolonging of di-liganded openings (Fig. 4E-G; Table S5). While this effect appears modest compared to that observed in the human AChR, the mutation also leads to changes in the single-channel burst behaviour, and thus its cumulative effect is more profound than indicated by open duration alone (Fig. S10). Regardless, it is important to note that the substitution is incorporated into both copies of AncDGE in this case, and thus it is unclear what the effect of the mutation is when installed in the δ versus ε-subunit positions alone. To begin teasing this apart, we performed an electrical fingerprinting experiment to measure open-channel lifetimes from AncDGE-containing AChRs incorporating only a single S276I substitution (Fig. 5). Mixing experiments paired mutation status (AncDGE versus AncDGE-S276I) with conductance (HC versus LC), to report on the number of incorporated AncDGE-S276I subunits. In this case, the S276I substitution was installed in the low-conductance AncDGE subunit (AncDGE-S276I_LC_). When AncDGE_HC_ and AncDGE-S276I_LC_ subunits are co-transfected, a distribution of single-channel amplitudes is observed within each patch. As expected, the amplitudes segregate into three distinct classes (Fig. 5A,B), where openings with the highest and lowest amplitudes derive from channels with two AncDGE_HC_ or two AncDGE-S276I_LC_ subunits, respectively. Openings falling into the intermediate amplitude class presumably originate from channels incorporating a single AncDGE_HC_ subunit, as well as a single AncDGE-S276I_LC_ subunit (Fig. 5B, bottom).

**Figure 5.**
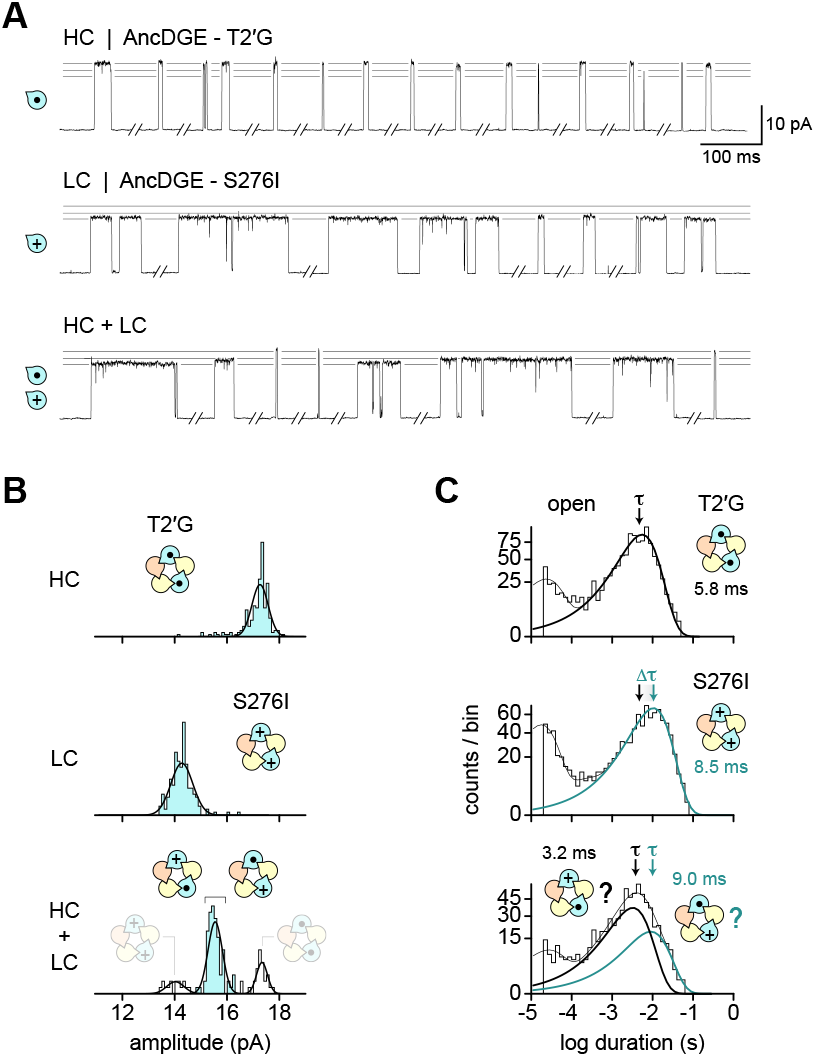
Positional dependence of AncDGE mutations. **(A)** Single-channel traces (1 µM acetylcholine; –120 mV) from cells transfected with cDNAs encoding human α, β, and either HC *(top)*, LC *(middle)*, or HC + LC *(bottom)* AncDGE subunits, where AncDGE_LC_ carries the S276I substitution (plus sign), and the AncDGE_HC_ carries a T2*′*G conductance mutation (black dot). Traces are shown filtered at 1 kHz, but were analyzed at 10 kHz. **(B)** Event-based amplitude histograms for recordings in panel A. Amplitudes were pooled from a minimum of three patches. For the HC + LC experiment, the middle amplitude class (cyan) represents AChRs carrying a single AncDGE-S276I substitution. **(C)** Open-duration histograms for AChRs in the HC and LC experiments, as well as openings from the intermediate amplitude class in the HC + LC experiment. Mean time constants (τ) for the di-liganded open components are indicated, with black being ‘wild-type’ and cyan being the serine-to-isoleucine substitution (S276I). For the HC + LC experiment, di-liganded openings are best-fit by two exponential components instead of one, as in the HC and LC experiments (log-likelihood ratio test, *p*-value < 0.001; see Fig. S11

We then measured open-channel lifetimes for each of the three amplitude classes (Fig. 5C). Open-duration histograms from AncDGE-containing AChRs incorporating two copies of ‘wild-type’ (AncDGE_HC_), or two copies of the mutant (AncDGE-S276I_LC_), each show a single-class of di-liganded openings, whose lifetimes correspondingly matched that of ‘wild-type’ or mutant AncDGE-containing AChRs (Fig. 5C; top and middle). In contrast, AncDGE-containing AChRs harbouring a single AncDGE_HC_, and a single mutant AncDGE-S276I_LC_, exhibit two types of diliganded openings, whose lifetimes match that of either ‘wild-type’ or mutant AncDGE-containing AChRs (Fig. 5C, bottom; Fig. S11). Assuming that the AncDGE_HC_ and AncDGE-S276I_LC_ subunits are stochastically incorporated, the intermediate amplitude class should be a mixture of channels, with some carrying the S276I substitution in the δ-position, and others in the ε-position. The observation that channels incorporating a single S276I substitution exhibit either the full effect of the mutation, or no effect at all, is consistent with the substitution having an effect in one position (presumably the ε-position), but not in the other.

To pinpoint where the AncDGE-S276I substitution exerts its effect, we selectively incorporated a single AncDGE subunit into the pentamer, at either the δ or the ε-position, and assessed the effect of the mutation in each position individually. Although AncDGE can substitute for both the δ and ε-subunits in AncDGE-containing AChRs, it does not have to replace both subunits simultaneously. Adding excess δ-subunit cDNA during transfection produces AChRs containing a δ-subunit as well as a single AncDGE subunit (Fig. 6A) Likewise, adding excess ε-subunit cDNA leads to AChRs incorporating an ε-subunit, and a single AncDGE subunit (Fig. 6C). Both the δ and ε-subunits are entrenched in specific positions, and thus their presence in the pentamer serves to direct AncDGE into a predetermined location. When co-transfected with δ, AncDGE is restricted to the ε-position (AncDGE_ε_), whereas co-transfecting with ε limits AncDGE to the δ-position (AncDGE_δ_). Fortuitously, AChRs incorporating a single AncDGE subunit can be dis-tinguished from those carrying two AncDGE subunits by either their single-channel kinetics (e.g., AncDGE_ε_; Fig. S12 or amplitudes (e.g., AncDGE_δ_; Fig. S12D)

**Figure 6.**
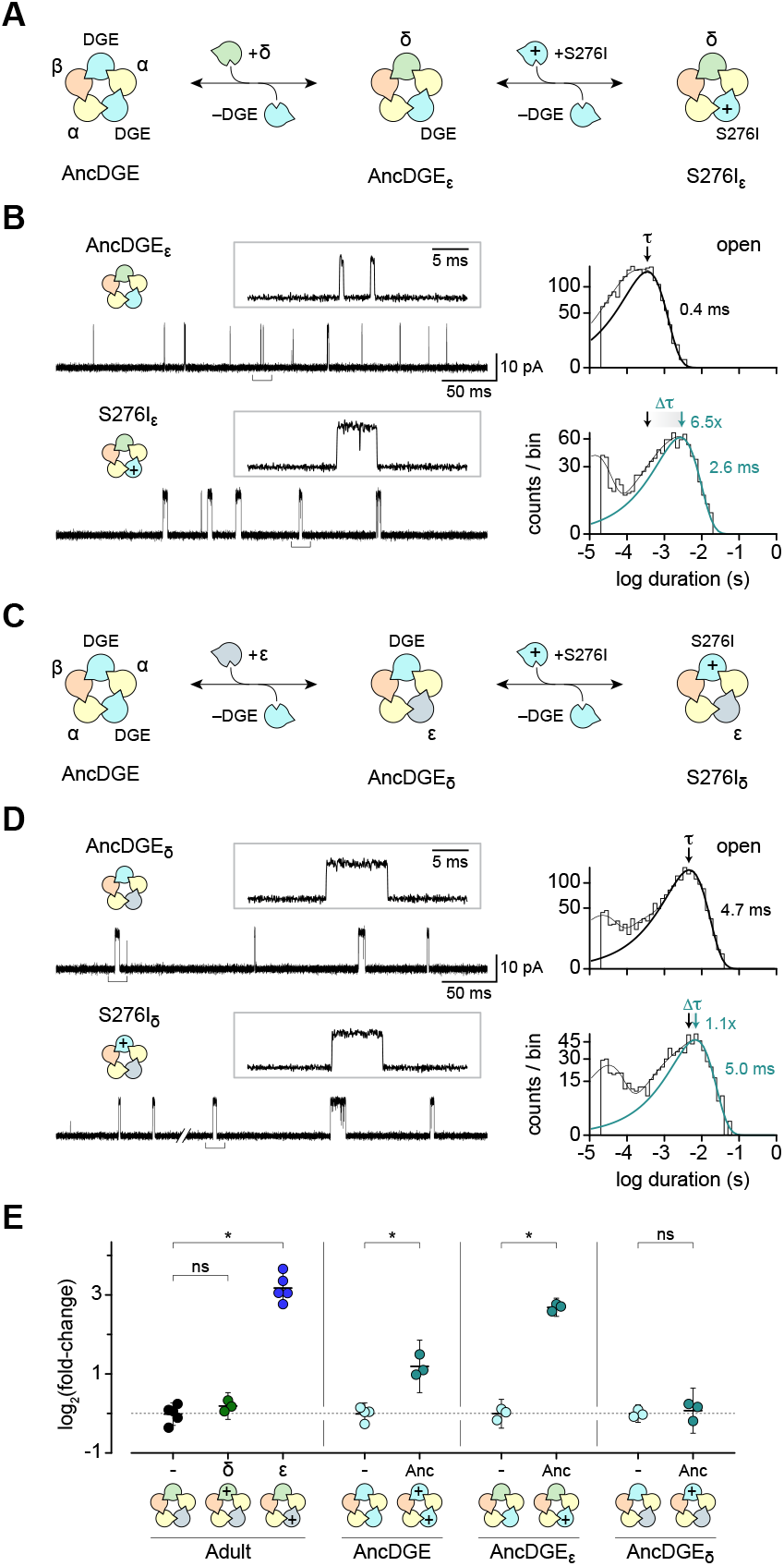
Single-subunit swap experiments delineate the positional dependence of AncDGE mutations. **(A**,**C)** Co-transfecting with δ-subunit (or ε-subunit) cDNA limits AncDGE incorporation to the ε-position (or δ-position). **(B**,**D)** Single-channel activity of AChRs incorporating a single AncDGE or AncDGE-S276I subunit in either the ε-position *(panel B)* or δ-position *(panel D)*. Open-duration histograms are shown with di-liganded exponential components bolded: AncDGE, black; AncDGE-S276I, cyan. **(E)** Effect of serine-to-isoleucine substitutions on di-liganded lifetimes. Data is presented as fold-change versus the appropriate non-mutant/’wild-type’ background. Mean ± 95 % confidence intervals are shown alongside individual replicates (n = 3-5 patches). *p*-values were calculated using a one-way ANOVA with Šídák multiple comparison correction. *p*-value: ‘^*^‘ < 0.001; ‘ns’, not significant, *p*-value > 0.05.

To better understand the positional dependence of the AncDGE-S276I substitution, we leveraged these single-subunit swaps to selectively mutate AncDGE in only the δ or ε-position (Fig. 6; Table S5). When AncDGE is restricted to the δ-position (AncDGE_δ_), the mutation has a negligible effect on di-liganded open-channel lifetimes (Fig. 6D) Conversely, when AncDGE is restricted to the ε-position (AncDGE_ε_), the same mutation prolongs openings 6.5-fold (Fig. 6B) Given that the same mutation is made in the AncDGE background in both cases, these results demonstrate that it is the position of the mutated subunit within the pentamer that determines if it has an effect. This position-dependence suggests that the observed δ or ε-subunit-dependent effects of the mutation in human AChRs are likewise driven by position, rather than by differences in the amino acid sequences of the δ and ε-subunits.

The positional dependence of the AncDGE-S276I substitution suggests that specific intersubunit interactions influence open-state stability. As mentioned previously, the two AncDGE subunits differ in whether their (+)-side interacts with the (–)-side of an α-subunit, or the (–)-side of the β-subunit (Fig. 4E) Given that the mutation stabilizes openings exclusively from the ε-position, interactions with the (–)-side of an α-subunit appear to be necessary. The (+)-side of the human β-subunit also interacts with the (–)-side of an α-subunit, and indeed installing the homologous substitution into the human β-subunit (βS279I) similarly prolongs di-liganded openings ∼9-fold (Fig. S9 Table S5). Together, these data suggest that intersubunit interactions across the two α(–) interfaces are important for open-state stability.

To verify that the increase in di-liganded open-state stability stems from interactions at the β(+) and α(–) interface, we took advatange of a previously reconstructed ancestral β-subunit (β_Anc_) (Prinston et al., 2017 which is able to occupy both the β and δ-positions in hybrid human/β_Anc_-containing AChRs (Fig. 7A (Emlaw et al., 2021 β_Anc_ lacks residues that are important for agonist binding, which necessitated the use of a higher acetylcholine concentration (10 µM) to elicit openings. Recognizing that open-channel block by acetylcholine can lead to a shortening of apparent open-channel lifetimes at higher acetylcholine concentrations (Ogden and Colquhoun, 1985; Sine and Steinbach, 1984 we focused on how the mutation alters the characteristically low open-probability of β_Anc_-containing AChR single-channel bursts (Fig. 7C Table S6). Installing the equivalent substitution (β_Anc_S273I) into both β_Anc_ subunits increased burst open probability from 0.11 to 0.53 (Fig. 7C,D). To test whether the effect of the mutation was exerted from the β-position, we concatenated the two β_Anc_ subunits through a short polypeptide linker (Emlaw et al., 2021 and mutated each of the two subunits in the concatemer separately. Controls with the human AChR where the δ and β-subunits were concatened indicated that the concatemer is incorporated in a counterclockwise direction, with the first subunit specifying the canonical δ-position, and the second the β-position (i.e., δ–β; Fig. S13). Bursts originating from channels incorporating the mutation in the δ-position (β_Anc_S273I–β_Anc_) exhibit a low burst open probability (0.10), matching those of ‘wild-type’ β_Anc_-containing AChRs. In contrast, bursts from channels where the mutation is incorporated in the β-position (β_Anc_–β_Anc_S273I) display a high open probability (0.54), mirroring β_Anc_-containing AChRs where the β_Anc_S273I mutant is incorporated in both the δ and β-subunit positions (Fig. 7C). We further corroborated our interpretation by preparing a concatemer with the δ and β_Anc_ subunits (δ–β_Anc_; Fig. 7C). As expected, mutating the β_Anc_-subunit in this δ–β_Anc_ concatemer increased the burst open probability from 0.16 to 0.49 (Fig. 7C). Our finding that the mutant β_Anc_S273I-subunit only exerts its effect when occupying the β-position, and not the δ-position, suggests that, once again, the mutation stabilizes the open state specifically through its interactions with the (–)-side of the neighbouring α-subunit.

**Figure 7.**
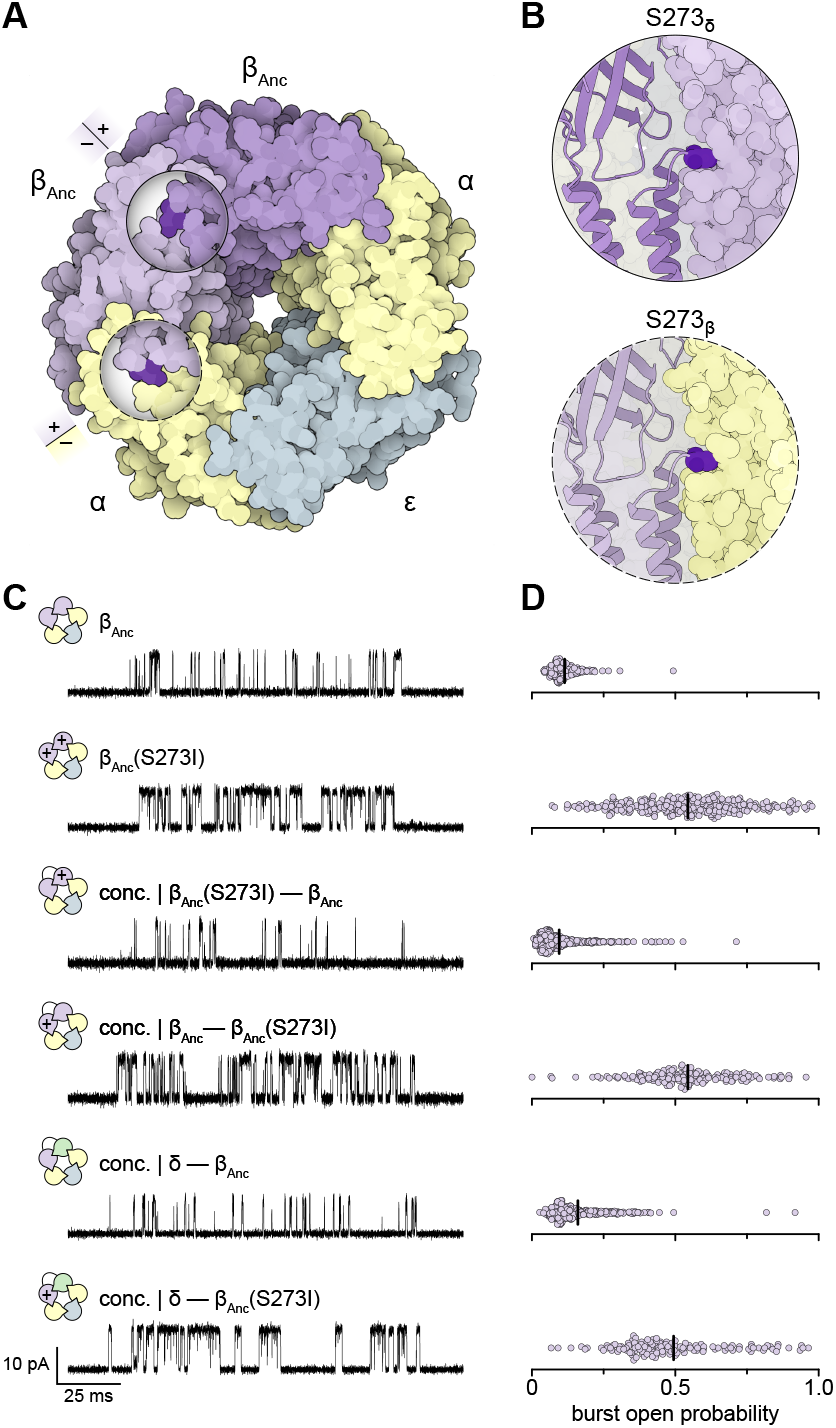
Positional dependence of β_Anc_ mutations. **(A)** Stoichiometry and arrangement of β_Anc_-containing AChRs. Circular cutaways show the S273I substitution (bold colouring) sitting at subunit interfaces. (+) and (–) sides of the interfaces are indicated. **(B)** Close-up views of the S273I substitution, as seen parallel to the membrane. **(C)** Representative single-channel bursts from AChRs incorporating zero to two β_Anc_ S273I substitutions. Subunit concatemers (‘conc.’ with connecting loop) were used to selectively mutate β_Anc_ in either the δ or β-position. Subunit rosettes illustrate stoichiometries and arrangements of each AChR, where subunits are coloured (α, yellow; β, orange; δ, green; β_Anc_, purple). Plus sign (+) denotes subunits bearing the S273I substitution. Single-channel bursts were elicited by 10 µM acetylcholine at –120 mV and are shown filtered at 10 kHz. **(D)** Distribution of burst open probabilities for corresponding AChRs in panel C. Open probabilities were pooled from three patches, with each circle representing the open probability of a single burst. Mean open probabilities are marked by the vertical bar.

## Discussion

With the aim of making an AChR with chemically identical agonist sites, we reconstructed an ancestral subunit representing the last common ancestor of the δ, γ, and ε-subunits. This ancestral subunit, which we call AncDGE, can substitute for the human δ and ε (or γ) subunits, and form functional AChRs when co-transfected with human α and β-subunits (Fig. 1). Using electrical finger-printing we show that AncDGE-containing AChRs have a 2α:1β:2AncDGE subunit stoichiometry, with AlphaFold predicting that the agonist sites are situated at the two chemically identical α(+)/AncDGE(–) interfaces (Fig. 2E,I). Consistent with this prediction, binding to AncDGE-containing AChRs is best fit by a ‘single-site’ model where the two agonist sites have the same apparent affinity for acetylcholine, and classic site-specific ligands (Fig. 2F-H). At low acetylcholine concentrations, single-channel openings from AncDGE-containing AChRs still exhibit two types of mono-liganded openings, indicating that acetylcholine binding to either of the two agonist sites alone does not stabilize the open-state equally (Fig. 3). Thus, despite being chemically identical, the two agonist sites in AncDGE-containing AChRs remain functionally inequivalent. By exploiting a human disease-causing mutation, we demonstrate that this functional inequivalence stems from asymmetric interactions across distinguishing subunit interfaces (Figs. 4-7). Together, our data indicate that the heteropentameric architecture of the AChR predisposes functional inequivalence of its two agonist binding sites, even in the absence of chemical differences. This principle may extend to other multi-subunit protein complexes, suggesting a generalizable mechanism by which oligomeric architecture shapes function across protein families.

This study adds to a growing body of work employing ancestral sequence reconstruction to explore aspects of AChR structure, function, and evolution (Lipovsek et al., 2014; Tessier et al., 2023, 2022, 2017; Emlaw et al., 2021, 2020; Prinston et al., 2017 Conceptually, the AncDGE-containing AChR, in which AncDGE replaces both the ε and δ-subunits, can be thought of as a ‘mutant’ human AChR containing 434 individual mutations, which represent 18.6 % (or 434 of 2,321 total amino acids) of the amino acid sequence of the adult human muscle-type AChR. Despite these extensive differences, the single-channel activity of AncDGE-containing and human AChRs is remarkably similar. In particular, the equilibrium constant describing apparent di-liganded gating (see ‘Θ_2_‘ in Table S3) is virtually indistinguishable between the two types of channels. While it is prudent to be cautious about the evolutionary implications of our experiments, especially since they involve hybrid ancestral/human AChRs, as opposed to ‘fully ancestral’ AChRs, the observed similarity in Θ_2_ suggests that the amino acid determinants of di-liganded gating, and in this case acetylcholine efficacy, have been selected for and conserved across the complementary δ, γ, and ε-subunits.

Our results speak to the subunit-dependence of disease-causing mutations (Grosman et al., 2000; Shen et al., 2003, 2019, 2020), highlighting how structural context determines pathogenicity. It is well established that, when installed in different AChR subunits, analogous mutations can have different effects on AChR function. Indeed, the substitution studied in the present work, as well as the original disease-causing deletion (see Fig. S9C), prolongs individual AChR openings almost 10-fold when they occur in the ε-subunit, yet have no effect when installed in the δ-subunit. The ε and δ-subunits only share 46 % sequence identity, and occupy unique positions within the AChR heteropentamer. For ε and δ, their amino acid sequence and position in the heteropentamer is inextricably linked, making it impossible to disentangle the relative contribution of each property to the subunit-dependent effect of the mutation. To address this, we exploited the plasticity of AncDGE, which can occupy both the ε and δ-positions, allowing us to isolate the role of subunit position from the amino acid background. This revealed that the mutant AncDGE subunit only prolongs openings when it occupies the ε-position. We therefore conclude that it is not the amino acid background of the mutant subunit that is definitive, but rather its specific position within the heteropentamer. This underscores the importance of considering subunit position when interpreting genotype-phenotype associations of channelopathies.

Establishing the positional dependence of the disease-causing mutation allowed us to uncover the underappreciated importance of the α(–) interface to AChR open state stability. In all AChR subunits, the disease-causing mutation maps to the loop linking the 2^nd^ and 3^rd^ transmembrane helices (‘M2-M3 linker’), a structure known to influence AChR gating. In each case, the substituted residue is pinched between the extracellular and transmembrane domains at the (–)-side of the neighbouring subunit. In ε and β, this is the (–)-side of one of the two α-subunits, while in δ it is the (–)-side of the lone β-subunit. Given that the mutation has an effect when installed in either ε or β, but not in δ, interactions with the α(–)-side appear essential. Precisely how the mutation causes an increase in open state stability through interactions at the α(–) interface is unclear. In both ε and β, the substituted residue interacts with key gating elements on the α-subunit: [1] the base of the β8-β9 hairpin structure constituting ‘loop C’; and [2] the linker connecting β10 and the first transmembrane helix (‘pre-M1 linker’) (Lee and Sine, 2005; Lee et al., 2009; Purohit and Auerbach, 2007; Mukhtasimova and Sine, 2013). Loop C from each α-subunit undergoes marked conformational change upon agonist binding, where it caps its respective agonist-binding site, trapping bound agonist and priming the AChR for opening (Zarkadas et al., 2022; Thompson et al., 2025). Meanwhile, translation of the α-subunit pre-M1 linker towards its neighbouring ε and β-subunit occurs concurrently with movements of the αM2-M3 linker, which open the channel pore. We also note that the ε-subunit effectively bridges the two AChR agonist-binding sites by [1] directly contributing residues to the α–ε site, and [2] indirectly, by interacting with the base of the loop C that caps the α–δ site. Consequently, the ε-subunit is uniquely poised to influence AChR activation through both agonist sites.

Phylogenetic analysis of vertebrate AChR subunits suggests that they evolved from an ancestral subunit that could form homopentamers (Fig. 1B; Fig. S1, S2). The current AChR subunit repertoire is thus the product of a series of gene duplication events. One of the earliest duplications led to two daughter genes that encoded for the ancestors of modern-day principle (+) and complementary (–) subunits. These ancestral subunits eventually lost the ability to form homopentamers, necessitating their coassembly as heteropentamers composed of both subunit types. Because pentamers contain an odd number of subunits, this initial heteromerization event was pivotal, disrupting the channel’s C5 symmetry. Subsequent duplications led to increasingly complex heteropentamers formed from three, and as in the muscle-type AChR, eventually four, different subunit types. A consequence of these latter duplications was that, as the amino acid sequences of the complementary subunits diverged, the chemistry of the AChR agonist sites became increasingly distinct. Our finding that the agonist sites remain functionally inequivalent in an AChR with chemically identical binding sites suggests that although divergence of the complementary subunits made specialization of the agonist sites possible, it was not necessary in order to establish their functional inequivalence. Instead, functional inequivalence of the agonist sites may have predated the divergence of the complementary subunits, and can stem solely from the pentameric architecture of a heteromeric AChR. More broadly, these findings highlight how the architecture of a heteromeric assembly can dictate functional inequivalence independent of sequence divergence, thereby shaping function and evolution in multimeric protein complexes.

## Methods

### Materials

Unless stated otherwise, general chemicals were purchased from Sigma-Aldrich or Fisher Scientific. TLC grade (>99 % purity) acetylcholine chloride was used for all electrophysiology experiments and purchased from Sigma-Aldrich. *d* -Tubocurarine was purchased in the form of (+)-Tubocurarine hydrochloride pentahydrate (CAS: 6989-98-6) from Sigma-Aldrich (Cat. No: T2379). [^125^I]-α-Bungarotoxin ([^125^I]-α-Btx) was purchased from Revvity (formerly PerkinElmer). All forms of synthetic DNA (e.g., primers and gene blocks) were supplied by Integrated DNA Technologies or Thermo-Fisher Scientific. All protein structure images were generated with either Illustrate (Goodsell et al., 2019) or ChimeraX (Meng et al., 2023).

### Ancestral Sequence Reconstruction

Ancestral protein reconstruction uses a multiple sequence alignment, molecular phylogeny, and a best-fit model of amino acid evolution to reconstruct probabilistic sequences of ancient proteins. Individual multiple sequence alignments were prepared for non-α AChR subunit paralogs (β1, β2, β4, δ, γ, and ε) and later combined. Sequences were recovered using NCBI Protein BLAST (Altschul et al., 1990) (https://blast.ncbi.nlm.nih.gov/Blast.cgi) using the appropriate human sequence as query. Sequences were aligned using MUSCLE (Edgar, 2004), and those with long insertions or deletions, or those containing unknown amino acids were removed. Molecular phylogenies were prepared from these multiple sequence alignments using PhyML-SMS (Guindon et al., 2010), with support for the inferred phylogeny based on SH-like approximate-likelihood (aLRT SH-like) ratio tests (Anisimova and Gascuel, 2006). Alignments were iteratively pruned until the topology of the resulting phylogenetic tree matched the accepted species phylogeny curated in the Open Tree of Life (Hinchliff et al., 2015; Michonneau et al., 2016). Individual ortholog alignments were prepared separately before being combined into a single ‘complementary paralog’ alignment and phylogeny. Additional pruning ensured that relationships between orthologs (Hinchliff et al., 2015), and paralogs (Pedersen et al., 2019), matched established topologies.

Reconstructed sequences, particularly those at the most ancestral nodes, are influenced by the outgroups used to root the tree. This effect diminishes the farther the reconstructed sequence is away from the root node. To minimize outgroup influence, and maximize the information feeding into our complementary subunit paralog tree, we created a second minimal phylogeny containing three to four sequences from all human AChR paralogs and rooted this tree using two 5-HT_3A_ sequences as outgroups. Sequences were again chosen such that the relationships between orthologs and paralogs in the resulting tree matched the expected topology. Portions of the cytoplasmic domain which did not share clear homology to the complementary subunits, and indels, which would introduce gaps into the complementary subunit alignment, were manually removed from the alignment.

We then combined this expanded paralog alignment with the complementary subunit alignment and realigned using MUSCLE (Edgar, 2004). The phylogenetic tree generated from this MUSCLE alignment was then used to inform our final alignment which used the phylogeny-aware algorithm PRANK (Löytynoja and Goldman, 2005). Our final phylogenetic tree, derived from this PRANK alignment, was mostly congruent with the expected topology, with discrepancies restricted to the principal subunits forming heteromeric neuronal receptors (e.g., α3 and α6 subunits are swapped with α5 and β3 subunits) (Fig. S1). Nevertheless, given that these incongruencies are far removed from the ancestral node of interest, and isolated to a different part of the tree, they are expected to have minimal effect on the reconstructed sequence (Hanson-Smith et al., 2010).

The amino acid sequence of AncDGE was inferred using Lazarus (Finnigan et al., 2012). The multiple sequence alignment and phylogeny were prepared as described above, where the best-fit model of amino acid evolution was calculated from the alignment using Smart Model Selection in PhyML (Lefort et al., 2017). The reconstruction employed the Jones-Taylor-Thornton (JTT) evolutionary model. 5-HT_3A_ outgroup sequences were culled/dropped prior to reconstruction using the *ape* package in R (Paradis et al., 2004). Posterior probabilities for each site were calsculated within Lazarus, and maximum-likelihood ancestral sequences were reconstructed by successively concatenating the amino acid with the highest posterior probability at each site in the alignment. As a test, the maximumlikelihood amino acid sequence was then added to our alignment and a new phylogeny generated. The reconstructed sequence of AncDGE ‘fell’ at the expected ancestral node, indicating that it represented a plausible ancestral sequence. AncDGE’s amino acid sequence was reverse translated, and codon optimized for heterologous expression in human-derived cell lines. The signal peptide from the human AChR δ-subunit was appended to the 5^*′*^ end of the codon-optimized DNA sequence, and two successive stop codons (TGA and TAA) were added to the 3^*′*^ end. The resultant cDNA was ordered as a synthetic gene (Thermo-Fisher, GeneArt Strings DNA Fragments) and cloned into the pRBG4 plasmid using Gibson Assembly. Sanger sequencing confirmed the open reading frame.

Our initial attempt at reconstructing AncDGE was unsuccessful as this ancestral subunit was unable to substitute for either of the human δ or ε-subunits. No α-Btx binding sites were detected on the surface of cells when AncDGE replaced one of, or both, the δ or ε-subunits. We reasoned that this incompatibility may result from inclusion of ray-finned fishes (Actinopterygii; includes teleost fishes) in our phylogeny. Teleosts are potentially problematic since they have experienced a whole-genome duplication (Taylor et al., 2001; Hughes et al., 2018) and are presumed to have evolved at faster evolutionary rates (Amemiya et al., 2013; Venkatesh et al., 2014). Additionally, some non-teleost Actinopterygii have two β-subunits despite reportedly diverging prior to the teleost whole-genome duplication. For these reasons, we removed all Actinopterygii sequences and inferred a new phylogeny, which was used to reconstruct a new ancestral sequence (see Fig. S3D). This updated version of AncDGE (the one presented here) could successfully substitute for both the δ and ε-subunits (Fig. 1C).

Single-channel recordings of AncDGE-containing AChRs exhibit prolonged episodes of activation reminiscent of those observed with gain-of-function/slow-channel congenital myasthenic mutations. To facilitate kinetic analysis, we introduced a threonine-to-serine substitution at the 12^*′*^ position (T12^*′*^S) in AncDGE. The reverse substitution, serine-to-threonine, is a slow-channel mutation in the δ-subunit (Grosman and Auerbach, 2000), and so we reasoned that re-introducing the 12^*′*^S may restore wild-type-like function. Indeed, AChRs incorporating this T12^*′*^S mutant show burst-like behaviour, and exhibit a clear concentration dependence to burst open probability, reminiscent of human AChRs (Fig. 3; Fig. S6, S7). All references to AncDGE are thus in the context of this T12^*′*^S mutant background.

### Molecular biology

cDNAs encoding the human AChR subunits (α, β, δ, γ, and ε) cloned into the pRBG4 plasmid were provided by Steven M. Sine (Mayo clinic), while the ancestral β-subunit (β_Anc_) was reconstructed and then inserted into the same pRBG4 plasmid by Gibson assembly as described previously (Prinston et al., 2017). Inverse PCR was used to introduce point mutations into the different subunit cDNAs (Silva et al., 2017). All sequences were confirmed by Sanger sequencing.

### Mammalian cell culture

cDNAs encoding human AChR (α, β, δ, γ, and ε) and ancestral AncDGE or β_Anc_ subunits were transfected into BOSC 23 cells (Pear et al., 1993). Cells were maintained in Dulbecco’s modified Eagle’s medium (DMEM) containing 10 % (vol/vol) fetal bovine serum at 37 °C and 10 % CO_2_. Cells were plated onto 3.5 cm dishes for single-channel recordings or 6 cm dishes for α-Btx binding experiments and transfected by calcium phosphate precipitation. Transfections were terminated after up to 8 h by exchanging the media. Electrophysiology experiments were performed one or two days post-transfection, except for AncDGE recordings at 0.1 µM acetylcholine, which were performed up to four days post-transfection. Binding experiments were performed two days post transfection. Additional cDNA encoding green fluorescent protein was included for electrophysiology experiments to identify transfected cells.

### Synthesis of Waglerin-1

All solvents used for peptide synthesis were purchased from Fisher Scientific. Reagents were purchased at the highest available purity and used without further purification. Amino acid building blocks, N,N^*′*^-Diisopropylcarbodiimide (DIC), Oxyma Pure, and Piperazine were purchased from Oakwood Chemical (Estill, SC, USA). All amino acids were protected with an N-terminal Fmoc, and if necessary, sidechains carried standard acid-labile protecting groups. Peptide synthesis was accomplished on a CEM Liberty Blue (Charlotte, NC, USA) microwave assisted peptide synthesizer. Liquid chromatography-coupled mass spectrometry analysis was carried out using a Shimadzu LC20A highpressure liquid chromatography system (HPLC) coupled with a Shimadzu LCMS2020 single quad mass spectrometer with an electrospray ionization source. Peptide purification was carried out using an Agilent 1260 Infinity analytical HPLC.

Waglerin-1 is a 22 amino acid peptide (GGKPDLR-PCHPPCHYIPRPKPR) with a single disulfide joining the two cysteine residues (Cys9 and Cys13) (Schmidt et al., 1992). Walgerin-1 was synthesized using Fmoc solid-phase peptide synthesis (SPPS). Synthesis was initiated by loading Wang resin with Fmoc-Arg(Pbf)-OH. Wang resin (500 mg, 0.83 mmol g^−1^) was first swelled in Dimethylformamide (DMF, 5 mL) for 30 min at room temperature in a fritted peptide vessel. Fmoc-Arg(Pbf)-OH (10 equiv.) was dissolved in Dichloromethane (DCM, 10 mL) and cooled to 0 °C. DIC (5 equiv.) was added and the solution was stirred for 30 min at room temperature. The organic solvent was removed under reduced pressure and the residue was dissolved in minimal DMF and transferred to the resin. The resin solution was agitated at room temperature for 2 h. The solution was then flushed and washed with DCM (2 x 10 mL), DMF (2 x 10 mL) and DCM (2 x 10 mL) and the resin dried under high vacuum. Resin loading was determined to be 0.30 mmol g^−1^ using a standard Fmoc loading test (Gude et al., 2002). The loaded resin (0.333 mg, 0.1 mmol) was transferred to the reaction vessel and underwent microwave assisted SPPS. Amino acids (0.2 M in DMF) were successively coupled using DIC (0.5 M) and Oxyma Pure (1 M) at 75-90 °C for 125 s. Fmoc deprotection of each amino acid was performed prior to coupling using 5 % Piperazine in 9:1 N-Methyl-2-pyrrolidone (NMP):Ethanol solution at 75-90 °C for 65 s. A final deprotection of the N-terminal Fmoc afforded the linear on-resin Waglerin-1 peptide. The peptide was cleaved, and acid-labile protecting groups removed through incubation (2.5 h) at room temperature) with Trifluoroacetic acid (TFA):Triisopropyl silane (TIPS):H_2_O (95:2.5:2.5; 5 mL). The TFA solution was added dropwise to a cold Diethylether (Et_2_O) flask (50 mL), and the peptide was precipitated overnight at −20 °C. The precipitate was centrifuged to afford a brown solid, which was dissolved in water (10 mL), lyophilized, and confirmed through LC-MS analysis. The single disulfide bond between Cys9 and Cys13 spontaneously formed upon overnight incubation of the peptide in water (∼1 mg mL^−1^), pH adjusted to 8.3 with Tris, free base (Schmidt et al., 1992). Product formation was monitored through LC-MS analysis. Excess water was removed through lyophilization, and the peptide was purified by semi-preparative HPLC to afford purified Waglerin-1 (∼180 mg, >95 % purity, MS ESI+; observed *m/z* [M+5H]^+5^ = 505, [M+4H]^+4^ = 631, [M+3H]^+3^ = 841, [M+2H]^+2^ = 1261). Purified Waglerin-1 was lyophilized, resuspended in water, and immediately frozen at −20 °C until used in an experiment.

### Radioligand binding experiments

[^125^I]-labelled α-Btx (specific activity = 2200 Ci/mmol) was purchased from Revvity (formerly PerkinElmer). The active concentration was calculated on the day of each experiment assuming decay catastrophe (Schmidt, 1984; Loring et al., 1982). Under these conditions, the specific activity is presumed to remain constant while the concentration of α-Btx decreases following the half-life of the ^125^I radioisotope (59.6 days). [^125^I]-α-Btx was mixed with unlabelled (‘cold’) α-Btx to reach the desired specific activity for each experiment.

Cell surface expression of AChRs was measured through binding of [^125^I]-labelled α-Btx to transfected cells (Emlaw et al., 2021). Approximately 1 million BOSC 23 cells were plated onto 6 cm tissue culture dishes and transfected using calcium phosphate precipitation as described above. Cells were harvested two days post-transfection by gentle agitation in phosphate buffered saline (PBS; pH 7.4) with 5 mM ethylenediaminetetraacetic acid (EDTA). Cells were spun at 1000 rpm for 2 min, and cell pellets re-suspended in potassium Ringer’s solution containing [^125^I]-α-Btx (15 nM, specific activity of 30 Ci/mmol). Potassium Ringer’s solution contained (in mM) 140 KCl, 5.4 NaCl, 1.8 CaCl_2_, 1.7 MgCl_2_, 25 HEPES, and 30 mg L^−1^ bovine serum albumin, adjusted to pH 7.4 with KOH. Following a 1 h incubation at room temperature, cells were centrifuged for 1 min at 6000 rpm and deposited onto 25 mm GF/C microfiber filter discs (Whatman) using a Hoefer filtration manifold. Unbound toxin was removed by washing three times with Ringer’s solution or phosphate buffered saline. To minimize non-specific binding of [^125^I]-α-Btx, GF/C filter discs were pre-incubated in Ringer’s solution containing 1 % (w/v) bovine serum albumin for at least one hour and dried prior to use. All filtration steps were performed under 25-28 inHg (85-95 kPa). Filter discs were counted for two minutes in a Wizard2 1-Detector Gamma Counter (Perkin Elmer).

Kinetics of α-Btx association were measured by incubating transfected cells with specified concentrations of [^125^I]-α-Btx (specific activity of 10 to 25 Ci/mmol) for up to 1 h. Non-specific α-Btx binding was determined from cells transfected with cDNA encoding the β-subunit, which does not itself form α-Btx binding sites. Specific binding was calculated by subtracting non-specific binding from the total binding to human or AncDGE-containing AChRs. The fraction of sites bound was determined by dividing specific binding at each incubation time by the specific binding determined in the presence of a saturating concentration of α-Btx (50 nM for human and 100 nM for AncDGE). Association rate constants (*k* _on_) were calculated in GraphPad Prism (v9) by simultaneously fitting data collected from three α-Btx concentrations (4 nM, 10 nM, and 25 nM; Fig. S14A,B).

Binding of acetylcholine, *d* -tubocurarine, and Waglerin-1 were determined in competition against the initial rate of [^125^I]-α-Btx binding as described previously (Sine et al., 1994; Molles et al., 2002a). Following incubation with increasing concentrations of competing ligand (30 min for acetylcholine and *d* -tubocurarine, 1 h for Waglerin-1), [^125^I]-α-Btx was added and the cells were incubated for a further 30 min before filtering. The concentrations of [^125^I]-α-Btx (specific activity of 10 to 30 Ci/mmol) used were 10 nM and 21.5 nM for human and AncDGE-containing AChRs, respectively. These concentrations were chosen as they are sufficient to occupy half of the available sites in 30 min and share equal α-Btx association rates (Fig. S14C). After subtracting non-specific binding, fractional binding of the competing ligand was calculated as the ratio of specific binding in the presence of ligand to specific binding in the absence of ligand. For all three ligands tested, the maximum concentration used was sufficient to fully outcompete α-Btx binding. Reduction in α-Btx binding corresponds to fractional occupancy by the competing ligand. Data were fit to an equation for one-site or two-site competitive binding within GraphPad Prism (v9). Preference for one-vs-two site models were evaluated using the extra-sum-of-squares F-test (Table S2). All data were fit using a 95 % confidence interval.

### In-silico modelling

Models of *Torpedo*, human, and AncDGE-containing AChRs were generated using either AlphaFold2 multimer (Evans et al., 2022) (run on the COSMIC2 science gateway (Cianfrocco et al., 2017); https://cosmiccryoem.org/tools/alphafoldmultimer/) or AlphaFold3 (Abramson et al., 2024). Stoichiometries used for each prediction are as follows: 2α:1β:1δ:1γ for *Torpedo*, and fetal human; 2α:1β:1δ:1ε for adult human; and 2α:1β:2AncDGE for AncDGE-containing AChRs. A homology model of β_Anc_ AChRs was generated in MODELLER 9.24 (Webb and Sali, 2016), using an AlphaFold2 multimer model of the human adult AChR as template. β_Anc_ was aligned to the δ and ε-subunits using the alignment 2d function in MODELLER (align2d). Alignment quality was assessed by comparing gap placements to a secondary alignment built from human AChR paralogs aligned with MUSCLE (Edgar, 2004). A minimum of 25 models were generated, and the best model was chosen based on its molpdf, DOPE, and GA341 MODELLER scores.

### Single-channel patch clamp recordings

Single-channel patch clamp recordings were performed as described previously (Mukhtasimova et al., 2016). Recordings from BOSC 23 cells transiently transfected with cDNAs encoding human adult or ancestral (AncDGE or β_Anc_) AChR subunits were obtained in the cell-attached configuration with a membrane potential of –120 mV and a temperature maintained between 19 and 22 °C. The pipette solution contained (in mM) 80 KF, 20 KCl, 40 K·aspartate, 2 MgCl_2_, 1 EGTA, and 10 HEPES, adjusted to pH 7.40 with KOH. Acetylcholine was added to the specified concentration in pipette solution, and stored at −80 °C. The external bath solution contained (again, in mM) 142 KCl, 5.4 NaCl, 0.2 CaCl_2_, and 10 HEPES, adjusted to pH 7.40 with KOH. Patch pipettes were fabricated from type 7052 or 8250 non-filamented glass (King Precision Glass), with inner and outer diameters of 1.15 mm and 1.65 mm, respectively, and coated with SYLGARD 184 (Corning). Single-channel currents from an Axopatch 200B patch clamp amplifier (Molecular Devices), with the gain set at 100 mV/pA and the internal Bessel filter at 100 kHz, were sampled at intervals of 1.0 µs using a BNC-2090 A/D converter with a National Instruments PCI 6111e acquisition card, and recorded to the hard disk of a PC by the program Acquire (Bruxton).

### Electrical fingerprinting

High and low conductance variants of each AChR subunit were generated by introducing substitutions at the 2^*′*^ position in the AChR pore known to alter single-channel amplitude (Villarroel et al., 1992; Imoto et al., 1991; Emlaw et al., 2021). Subunit stoichiometries were determined by transfecting high and low conductance variants alone, or in combination with the remaining subunits. Single-channel amplitudes were measured as described previously (Emlaw et al., 2020). Briefly, amplitudes were calculated from single-channel bursts as the difference between open and closed-channel components of an all-points histogram. Amplitudes were pooled from a minimum of three patches to generate event-based amplitude histograms. For subunit mixing experiments, which had multiple amplitude classes, means and standard deviations were calculated by fitting the data to a normal distribution using the *mixtools* package in R (Benaglia et al., 2009). Means and standard deviations (Table S1) were then used to generate Gaussian fits, which were overlaid onto each histogram. For each mixing experiment, at least one recording contained a representative burst for each amplitude class. All recordings were digitally filtered at 10 kHz inside TAC (v4.3.3; Bruxton). 30 µM acetylcholine was chosen for fingerprinting experiments in Fig. 2 because it elicits robust single-channel bursts of AncDGE-containing AChRs but does not lead to an appreciable reduction in single-channel amplitude via open-channel block. For the AncDGE-S276I mixing experiments presented in Fig. 5 a slightly different procedure was used. Openings were elicited by 1 µM acetylcholine instead of 30 µM, and amplitudes were measured for individual openings instead of bursts.

### Kinetic fitting of single-channel dwells

Kinetic fitting was performed using a strategy similar to that described previously (Lee and Sine, 2004). Single-channel recordings were analyzed using the program TAC (v4.3.3; Bruxton). After applying a 10 kHz digital Gaussian filter, unitary current amplitudes were calculated for several segments (n > 10) of each recording, and averaged to give a single amplitude value for each patch. This average amplitude was then used for the detection of opening and closing transitions using the half-amplitude threshold crossing criterion. Open duration and closed duration histograms were calculated inside TACFit (v4.3.3; Bruxton) as successive open-to-close and close-to-open transitions, respectively. Critical closed durations (τ_crit_), which delineate bursts originating from a single ion channel, were calculated from the closed duration histogram as the intersection of the slowest activation and fastest desensitization components (Sine et al., 1990; Colquhoun and Hawkes, 1982).

Kinetic information was extracted using the *scbursts* package in R (Drummond et al., 2019). Open and closed transitions were converted into dwell durations, corrected for the instrument risetime, and separated into bursts using the appropriate τ_crit_. Bursts containing fewer than three openings were omitted from further analysis. Open probabilities for individual bursts were calculated as the time spent open within a burst divided by the total duration of that burst. To get a kinetically homogenous dataset, bursts whose open probability was deemed an outlier (van der Loo, 2010), or which fell beyond two standard deviations from the mean (Venables and Ripley, 2002), were removed. To calculate the mean burst open probability at a given acetylcholine concentration (Fig. 3A), we first plotted the open probability of individual bursts within each patch, and fit the resulting histogram with a Gaussian distribution. The mean value of the Gaussian fit was then averaged across three independent patches per concentration. Global kinetic fitting was performed by simultaneously fitting triplicate patches across an acetylcholine concentration range using MIL (QuB) (Qin et al., 1996, 1997). Acetylcholine concentrations ranged from 3 µM to 180 µM for human, and from 6 µM to 100 µM for AncDGE, with dilutions spaced at approximately quarter log-unit intervals (i.e., 1.78-fold). Concentrations were chosen to provide coverage across the minimum to maximum range of burst open probabilities. A uniform dead time of 18.833 µs was applied to all data.

The sequence of single-channel dwells were fit to multiple schemes (Table S7). While several variations of the extended del Castillo & Katz scheme (Colquhoun and Sakmann, 1985) were assessed, our time resolution was not sufficient to include schemes with short-lived (i.e., ‘flipped’ or ‘primed’) intermediate states (Lape et al., 2008; Mukhtasimova et al., 2009; Thompson et al., 2025). Best fitting models were determined by log-likelihood ratio tests and the Schwarz information criterion (SIC) as described previously (Shelley et al., 2010; Mukhtasimova et al., 2016). Human adult AChRs were best fit to Scheme I. For AncDGE, Scheme II, which has mono-liganded openings removed, ranked slightly higher than Scheme I, but both schemes gave virtually identical rate constants (Table S3). We therefore used Scheme I for both human and AncDGE-containing AChRs to simplify comparisons.

### Low acetylcholine concentration recordings

Single-channel recordings were digitally filtered at 10 kHz and analyzed using TAC (v4.3.3; Bruxton). At low acetylcholine concentrations openings appear in isolation rather than as bursts of succesive openings and closings, and so amplitudes were estimated by manually fitting the baseline and open-channel current levels of individually resolved openings (n = 10-12). Average amplitudes from each patch were then used for the detection of opening and closing transitions using the half-amplitude threshold crossing criterion. Open-duration histograms were fit inside TACFit (v4.3.3; Bruxton) by a sum of exponentials. Likelihood-ratio tests were used to determine the appropriate number of components in each fit (e.g., Fig. S8) (Shelley et al., 2010; Landowne et al., 2013). Mean open-channel lifetimes were calculated as the average from three to six patches, except for AncDGE at 0.1 µM which, due to infrequent openings, was calculated after pooling openings from five separate patches. One-way ANOVA’s were performed in GraphPad Prism (v9) using a 95 % confidence interval and Tukey’s multiple-comparison correction.

## Supplementary Information

Supplementary Information is appended to this document, and includes Figures S1–S14 and Tables S1–S7.

## Acknowledgements

We thank Kathleen M. Gilmour for access to a γ-counter, David S. Goodsell for help with Illustrate (Goodsell et al., 2019), and Timothy Lynagh for helpful discussions. J.R.E. was supported by an NSERC CGS-D graduate scholarship (CGSD3-559665-2021). This work was funded by a Natural Sciences and Engineering Research Council of Canada Discovery Grant awarded to C.J.B.d.C. (RGPIN-2024-05272). This manuscript was prepared using a modified version of a L^A^T<SUB>E</SUB>X template kindly made available by Stephen Royle on GitHub (click here).

## Generative AI Declaration

In the latter stages of manuscript writing the authors used ChatGPT (OpenAI) and CoPilot (Microsoft) to suggest targeted text edits that could improve clarity. After engaging these tools, the authors reviewed, edited, and then incorporated select edits. The authors take full responsibility for the entire content and wording of the manuscript. No generative AI or AI-assisted tools were used in the experimental design and data interpretation, or in the preparation of the figures, images, and artwork associated with this work.

## Author Contributions

J.R.E. and C.J.B.d.C. conceptualized the project. J.R.E. performed ancestral reconstruction with input from C.J.B.d.C. Electrophysiology data was collected and analyzed by J.R.E., with help from C.J.G.T. (β_Anc_ and human concatemers) and M.T. (deletion mutants). J.R.E. performed α-Btx binding experiments. A.R.P. synthesized Waglerin-1 peptide under supervision from C.N.B.. C.J.B.d.C. generated AlphaFold models with assistance from J.R.E.. J.R.E. and C.J.B.d.C. wrote the paper, with input from all authors.

## Author Declaration

The authors declare that they have no conflicts of interest with the contents of this article.

## Supplementary Information

**Figure S1.**
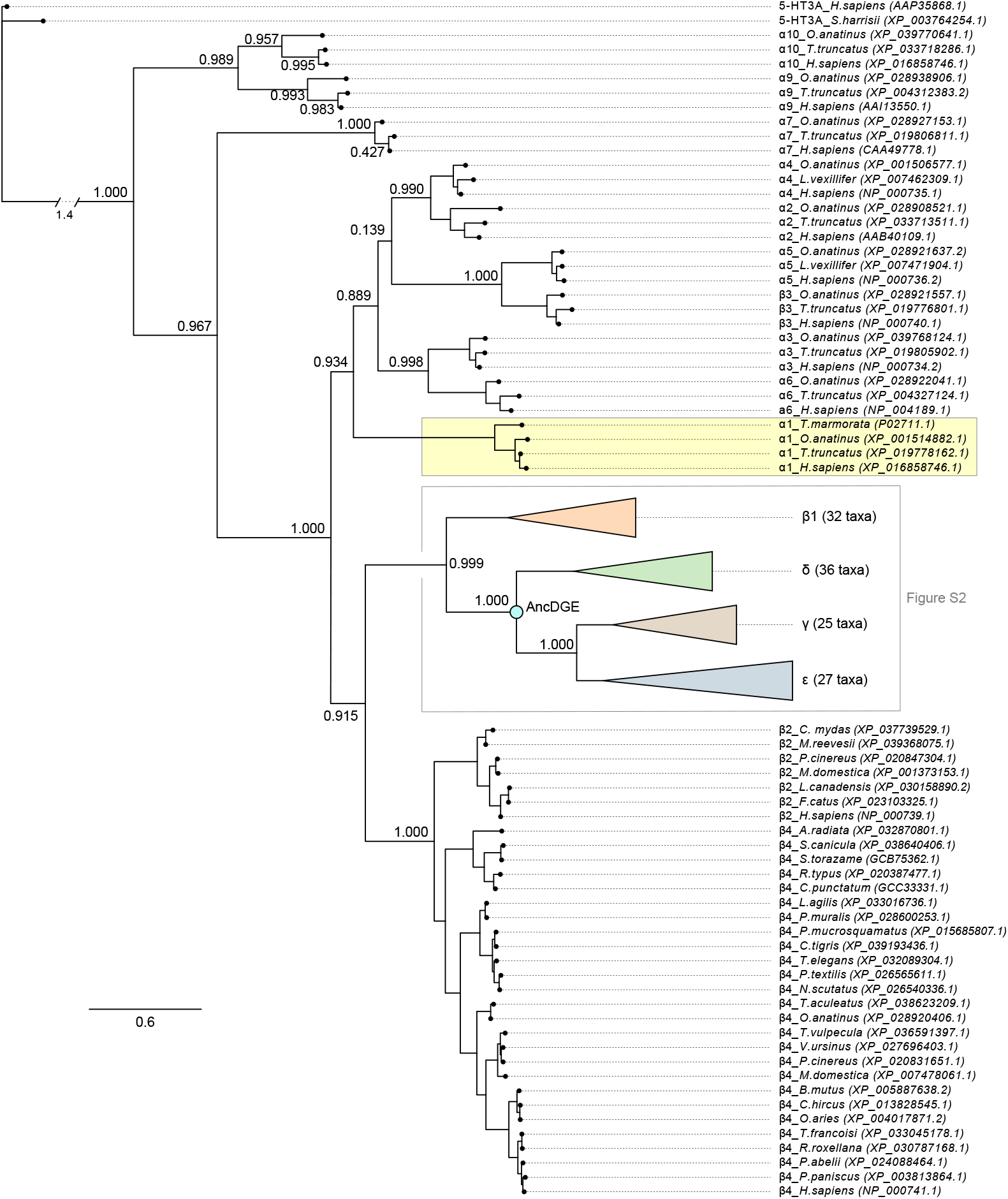
Ancestral reconstruction of AncDGE. AncDGE was reconstructed from a molecular phylogeny. A total of 186 sequences were included in the phylogenetic analysis. The two 5-HT_3A_ receptor sequences at the top are the outgroup sequences used to root the tree. Complementary muscle-type acetylcholine receptor subunits included in the phylogenetic analysis are collapsed for visualization (see Figure S2 for expansion). Labels on internal branches are approximate likelihood ratios with the nonparametric Shimodaira-Hasegawa (SH) correction, and the scale bar represents the average number of substitutions per site.

**Figure S2.**
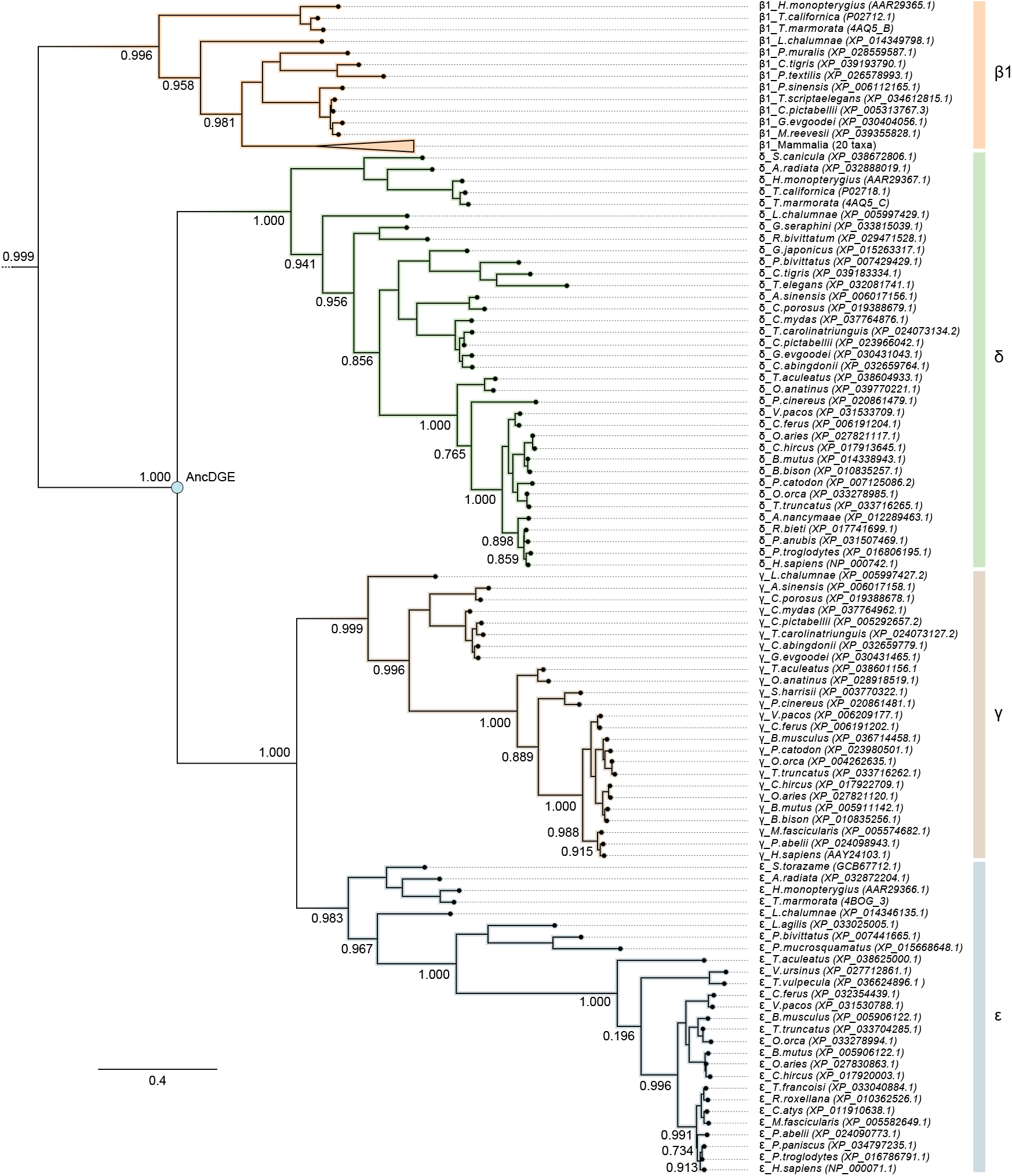
Non-α muscle-type acetylcholine receptor subunits used for ancestral sequence reconstruction. AncDGE was reconstructed from a molecular phylogeny where non-α muscle-type acetylcholine receptor subunits included in the analysis are shown, with mammals from the β1 subtree collapsed *(triangle)* to save space. Sequences corresponding to the 5-HT_3A_ outgroups, neuronal acetylcholine receptor subunits, and the α1-subunit can be found in Figure S1. Labels on internal branches are approximate likelihood ratios with the nonparametric Shimodaira-Hasegawa (SH) correction, and the scale bar is the average number of substitutions per site.

**Figure S3.**
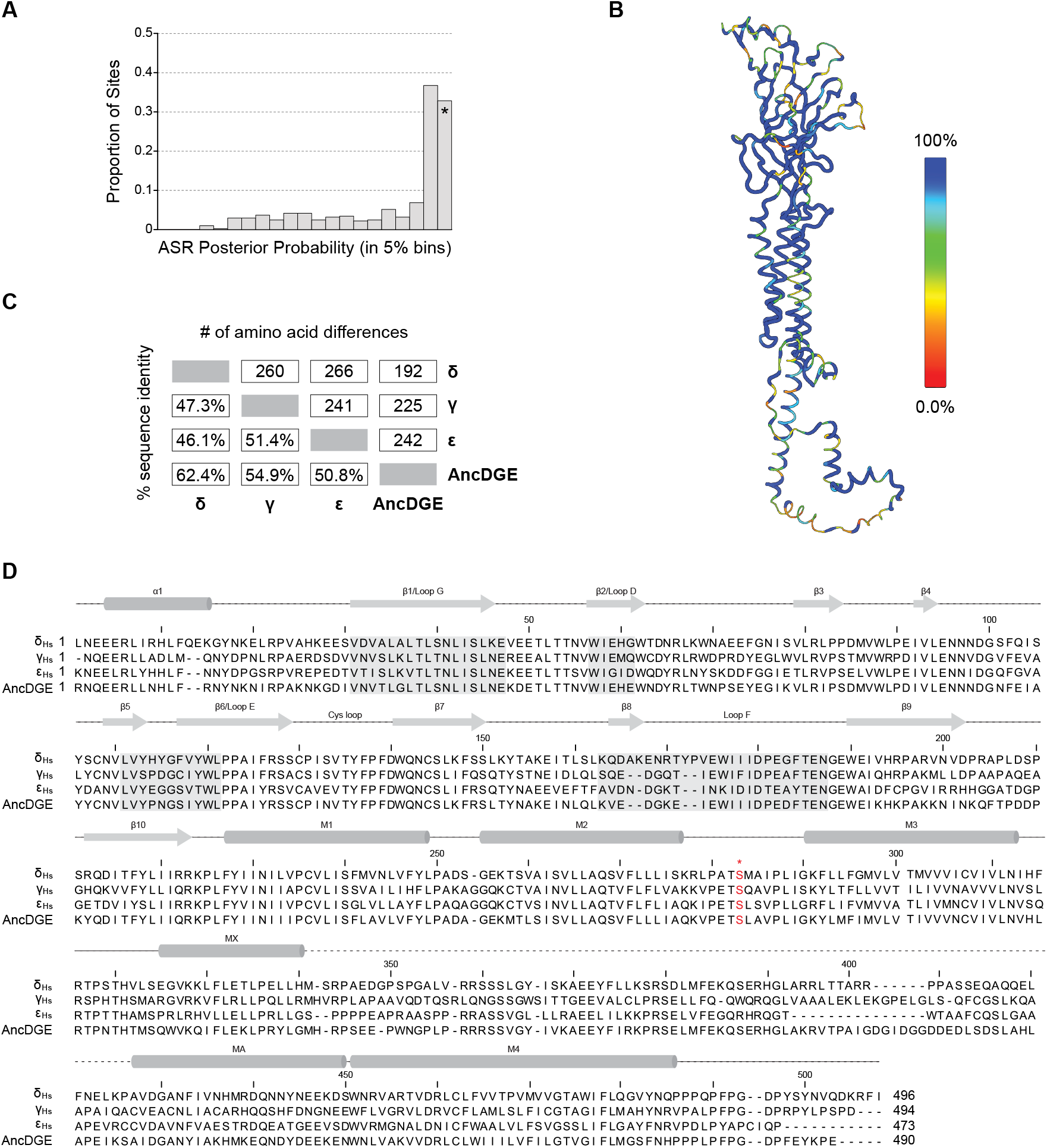
Bioinformatic analysis of AncDGE. **(A)** Posterior probability support for the reconstructed AncDGE sequence. Posterior probabilities for each position in the alignment are binned in 5 % intervals. The final bar (asterisk) corresponds to sites with 100 % posterior probability. **(B)** Posterior probabilities are mapped onto an AlphaFold2-multimer model of AncDGE. Spectrum bar on the right shows the relationship between posterior probability and colour. Areas coloured blue have higher posterior probabilities. **(C)** Matrix comparing sequence identity *(lower left)* and number of amino acid differences *(upper right)* between human δ, γ, ε, and AncDGE subunits. **(D)** Multiple sequence alignment of human (Hs) δ, γ, and ε-subunits and AncDGE. Secondary structure elements and important loops involved in agonist recognition (loops D, E, F, and G; boxed) as well as the eponymous Cys-loop are noted. The dashed line represents the cytoplasmic domain. The site of the εS278 substitution is noted in red with an asterisk.

**Figure S4.**
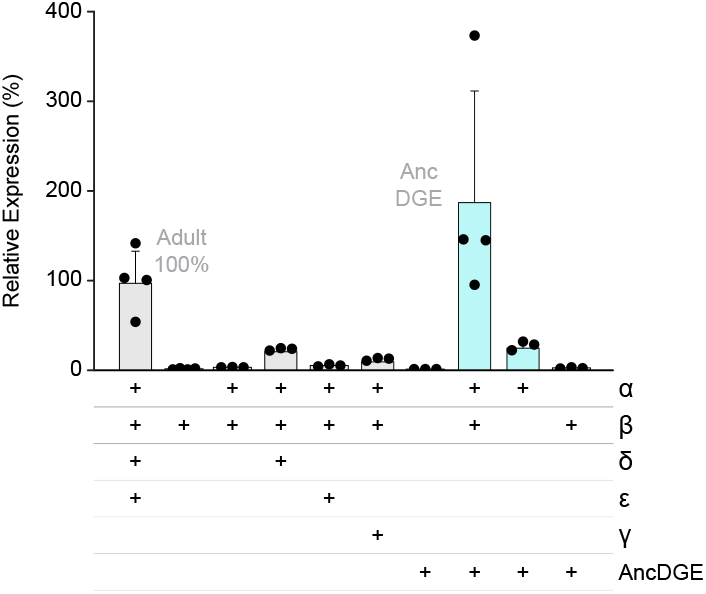
Cell-surface expression of AncDGE-containing acetylcholine receptors. [^125^I]-α-Btx binding to cells transfected with different combinations of human α, β, δ, γ, ε, and AncDGE cDNAs. Data was normalized to the mean expression of cells transfected with the full complement of adult human subunits (α, β, δ, and ε; 100 %), and is shown as mean plus standard deviation (n = 3-4 replicates) with individual replicates shown as black circles. The plus signs (+) in each column indicate the combination of cDNAs transfected.

**Figure S5.**
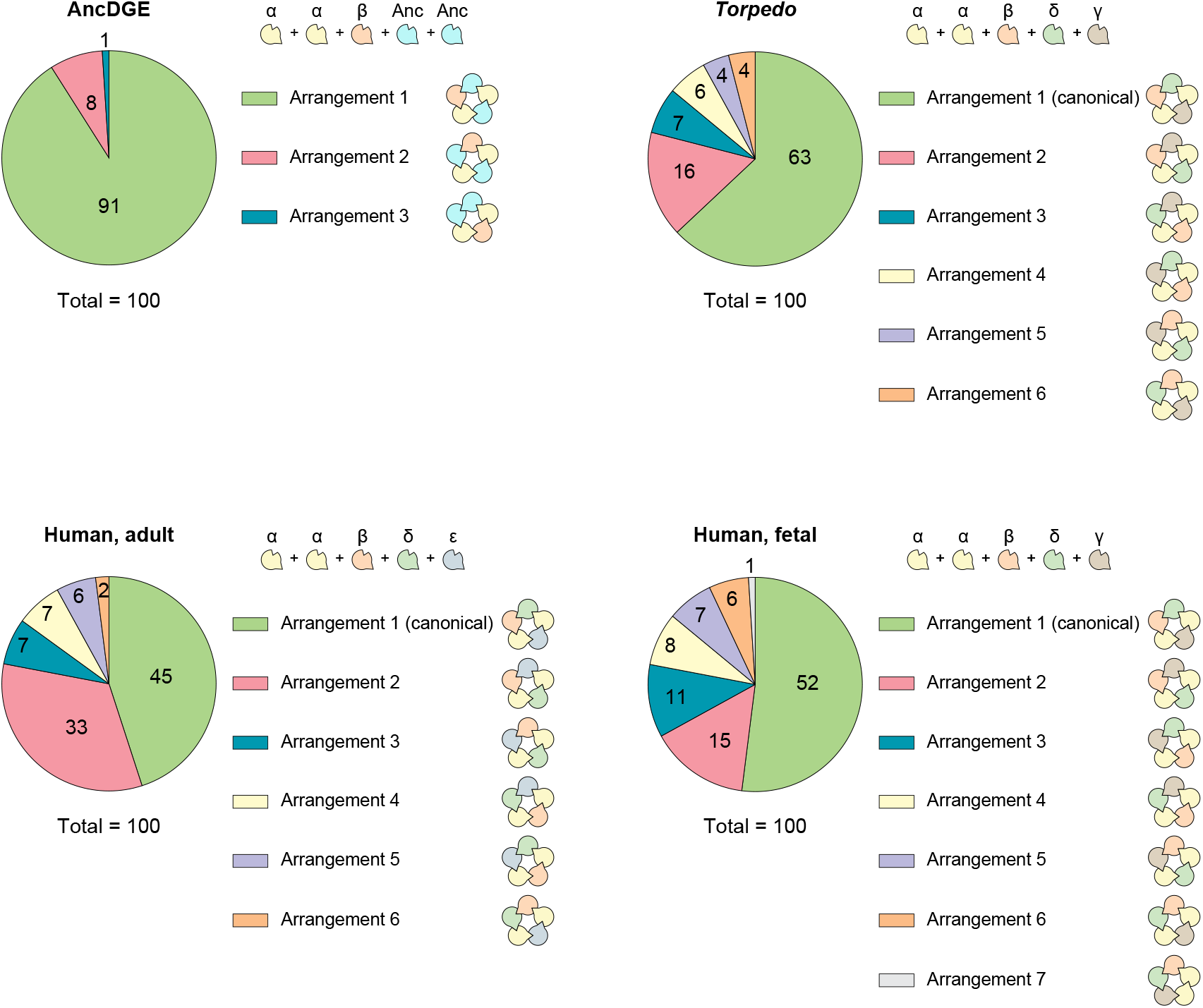
Predicted subunit arrangements of acetylcholine receptors. AlphaFold3 was used to predict the quaternary arrangement of AncDGE-containing AChRs *(top, left)*, as well as AChRs from *Torpedo (top, right)*, adult human *(bottom, left)* and fetal human *(bottom, right)*. One hundred predictions were generated for each set of sequences using known subunit stoichiometries for *Torpedo* and human AChRs (2α:1β:1δ:1γ/ε), and the experimentally determined stoichiometry for AncDGE-containing AChRs (2α:1β:2AncDGE). Different arrangements and the frequency in which they were predicted are shown. Subunits are colour-coded as follows: α, yellow; β, orange; δ, green; γ, brown; ε, blue; AncDGE, cyan.

**Figure S6.**
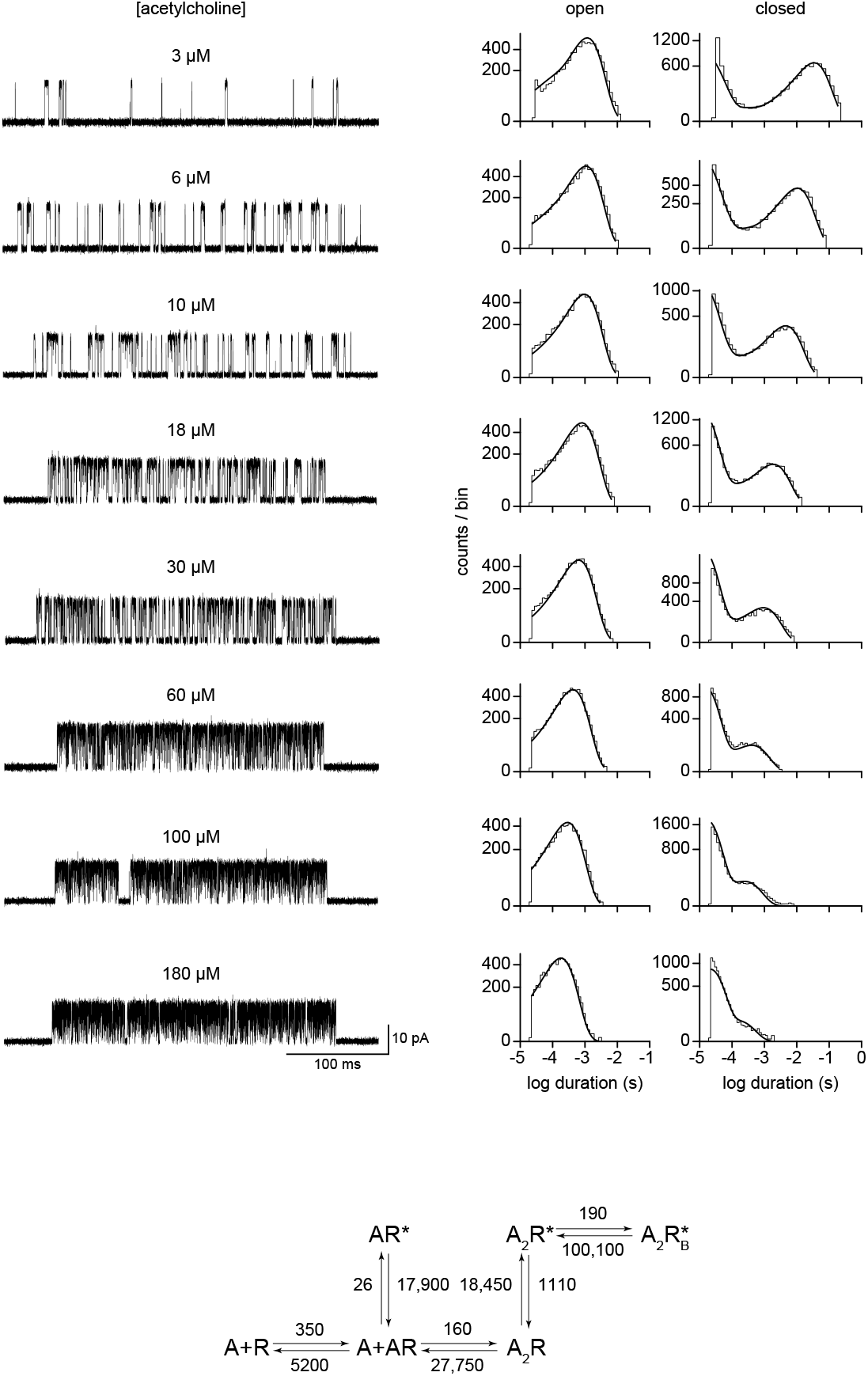
Global fitting of adult human AChRs. Openings and closings from single-channel bursts were fit to the extended del Castillo & Katz scheme (Scheme I; *bottom*). Dwells were obtained across a range of acetylcholine concentrations (3 µM to 180 µM) and simultaneously fit (in triplicate) to estimate transition rate constants. Representative single-channel bursts are shown alongside open and closed-duration histograms for each acetylcholine concentration. The resulting fit, from global fitting of the entire data set, is shown overlaid on the histograms. Transition rate constants (see Table S3) are shown above or below the appropriate arrow in the scheme (bottom). Single-channel recordings were obtained at –120 mV, and filtered to 10 kHz.

**Figure S7.**
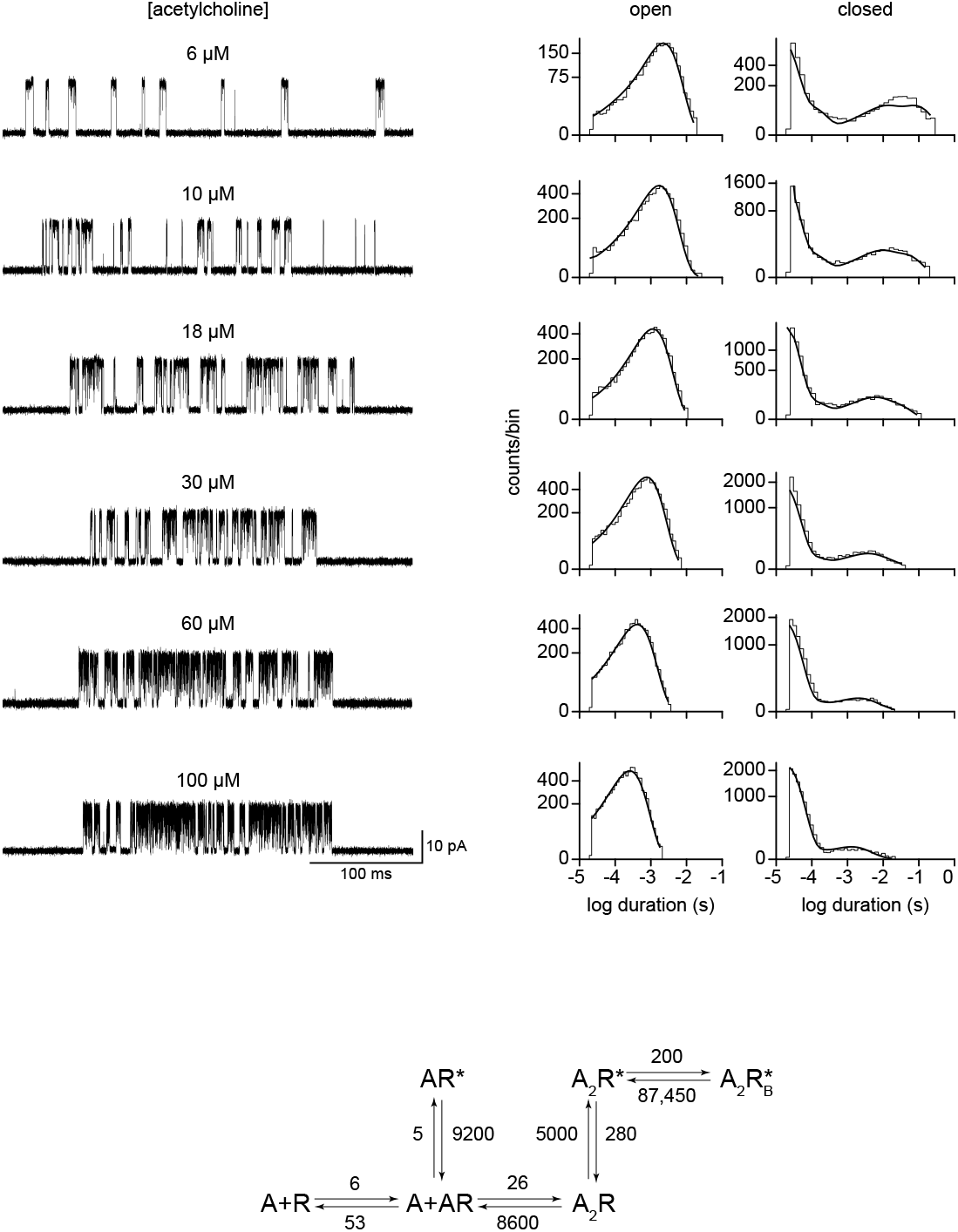
Global fitting of AncDGE-containing AChRs. Openings and closings from single-channel bursts were fit to the extended del Castillo & Katz scheme (Scheme I; *bottom*). Dwells were obtained across a range of acetylcholine concentrations (6 µM to 100 µM) and simultaneously fit (in triplicate) to estimate transition rate constants. Representative single-channel bursts are shown alongside open and closed-duration histograms for each acetylcholine concentration. The resulting fit, from global fitting of the entire data set, is shown overlaid on the histograms. Transition rate constants (see Table S3) are shown above or below the appropriate arrow in the scheme (bottom). Single-channel recordings were obtained at –120 mV, and filtered to 10 kHz.

**Figure S8.**
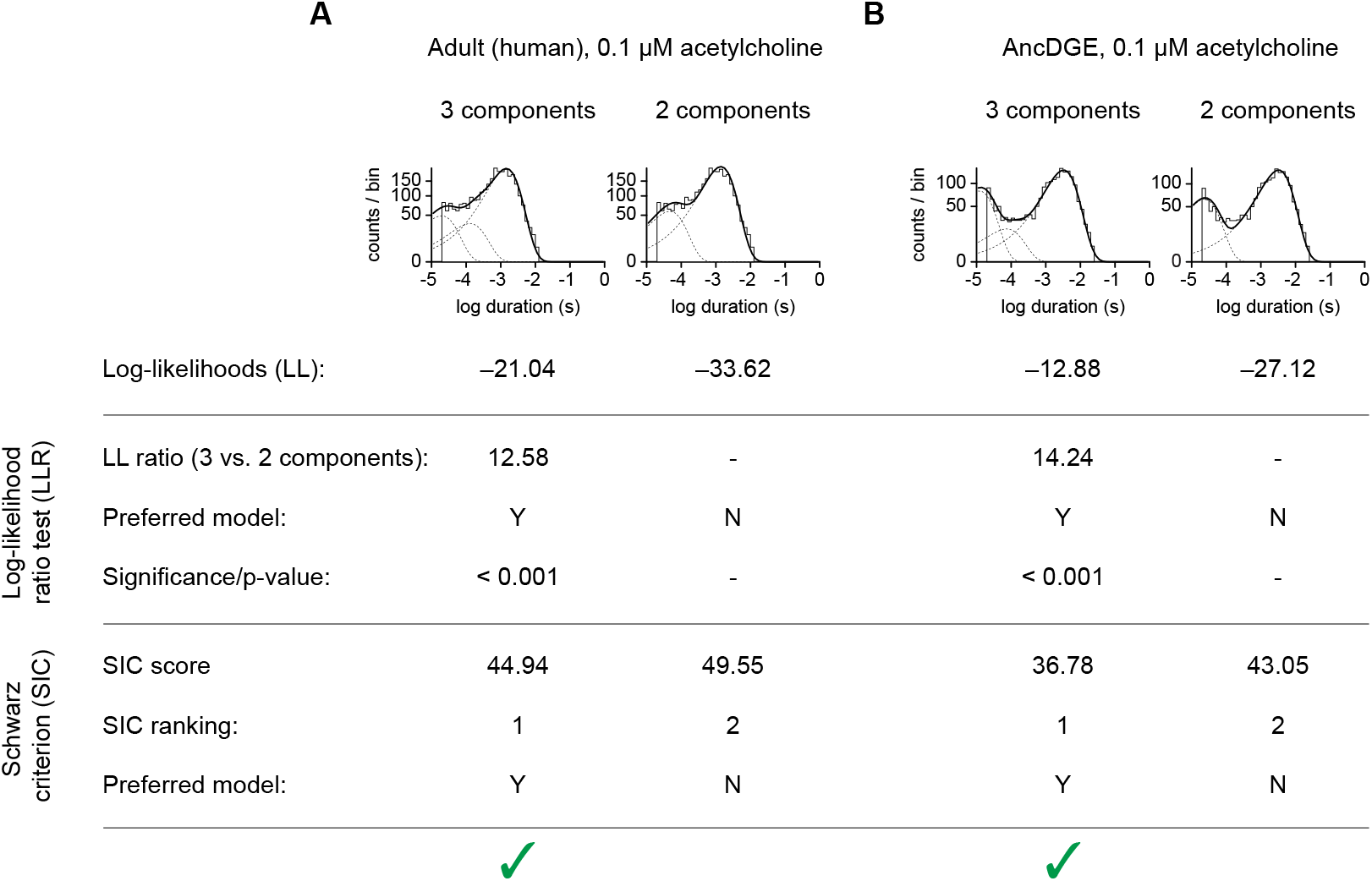
Fitting the correct number of components to open-duration histograms. Opening durations from **(A)** adult human, and **(B)** AncDGE-containing AChRs can be fit as the sum of two *(right)* or three *(left)* exponential components. The log-likelihood for both fits are provided below the histogram. Both the log-likelihood (LL) ratio test and Schwarz information criterion (SIC) scores indicate that the three component fits are most appropriate (green checkmarks). Log-likelihood ratios were calculated as LLR = LL(3 components) - LL(2 components). Signifiance was determined by comparing the LLR to the Chi-square table (2 degrees of freedom). SIC scores were calculated as SIC = -LL + (0.5*F* ) *×* ln(*N*), where *F* is the number of free parameters, and *N* is the number of observations. Single-channel openings were elicited by 0.1 µM acetylcholine.

**Figure S9.**
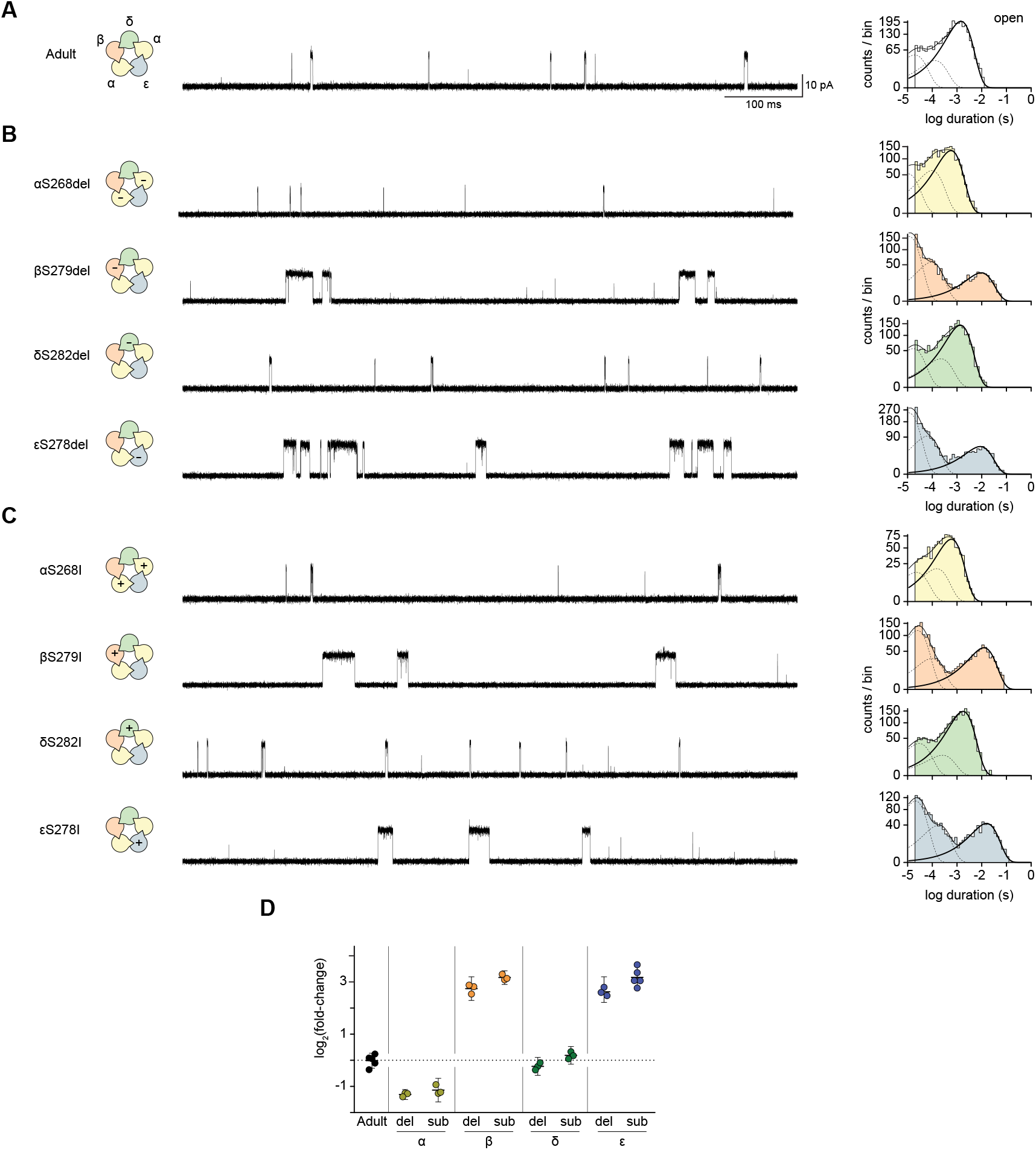
Comparing deletion and substitution mutations in human AChRs. Representative single-channel activity for **(A)** adult wild-type **(B)** deletion, and **(C)** substitution mutants. Single-channel recordings were obtained at 0.1 µM acetylcholine and –120 mV. Capacitor resets were removed for illustrative purposes. Scale bar in panel A (100 ms, 10 pA) applies to all traces. Open-duration histograms are coloured according to the mutated subunit: wild-type, white; α, yellow; β, orange; δ, green; ε, blue. Plus (+) and minus (–) signs indicate subunits with substitution (‘sub’) and deletion (‘del’) mutations, respectively. The longest exponential component in each histogram, which corresponds to the lifetime of di-liganded openings, is shown in bold and was used to determine the fold-change presented in panel D. **(D)** Fold-change in di-liganded open lifetime caused by each mutation is shown with 95 % confidence intervals (n = 3-5 patches).

**Figure S10.**
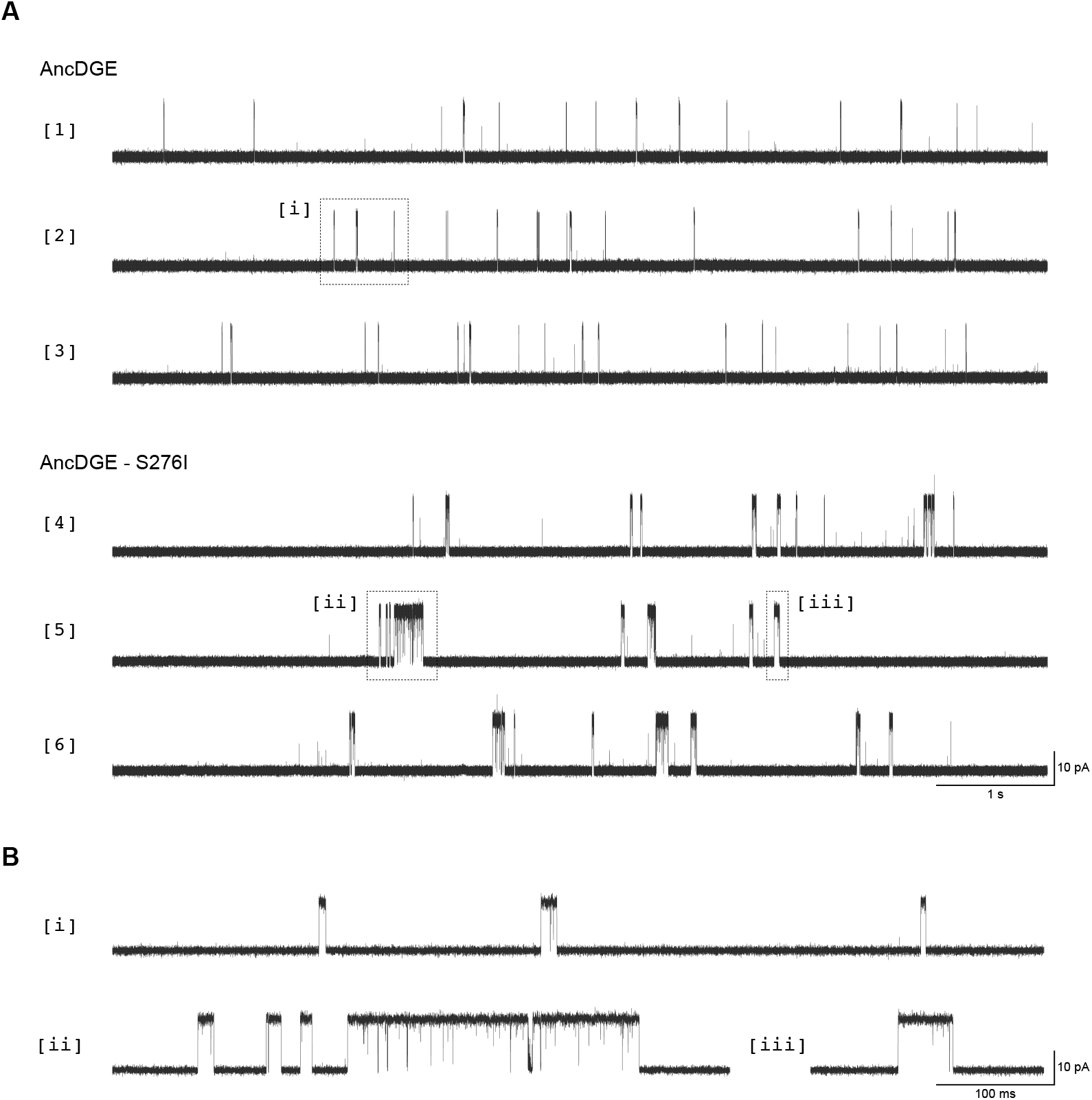
Single-channel activity of AncDGE-containing and AncDGE-S276I-containing AChRs. **(A)** Representative single-channel activity of AncDGE-containing (*top*; sweeps 1-3) and AncDGE-S276I-containing (*bottom*; sweeps 4-6) AChRs. Individual sweeps are different segments from the same recording. Activity was elicited by 1 µM acetylcholine, and recorded at –120 mV. Scale bar (1 s, 10 pA) applies to all sweeps in panel A. Dashed boxes [i], [ii], and [iii] indicate openings expanded in panel B. **(B)** Zoom-ins of selected openings from panel A. All data is shown digitally filtered at 10 kHz.

**Figure S11.**
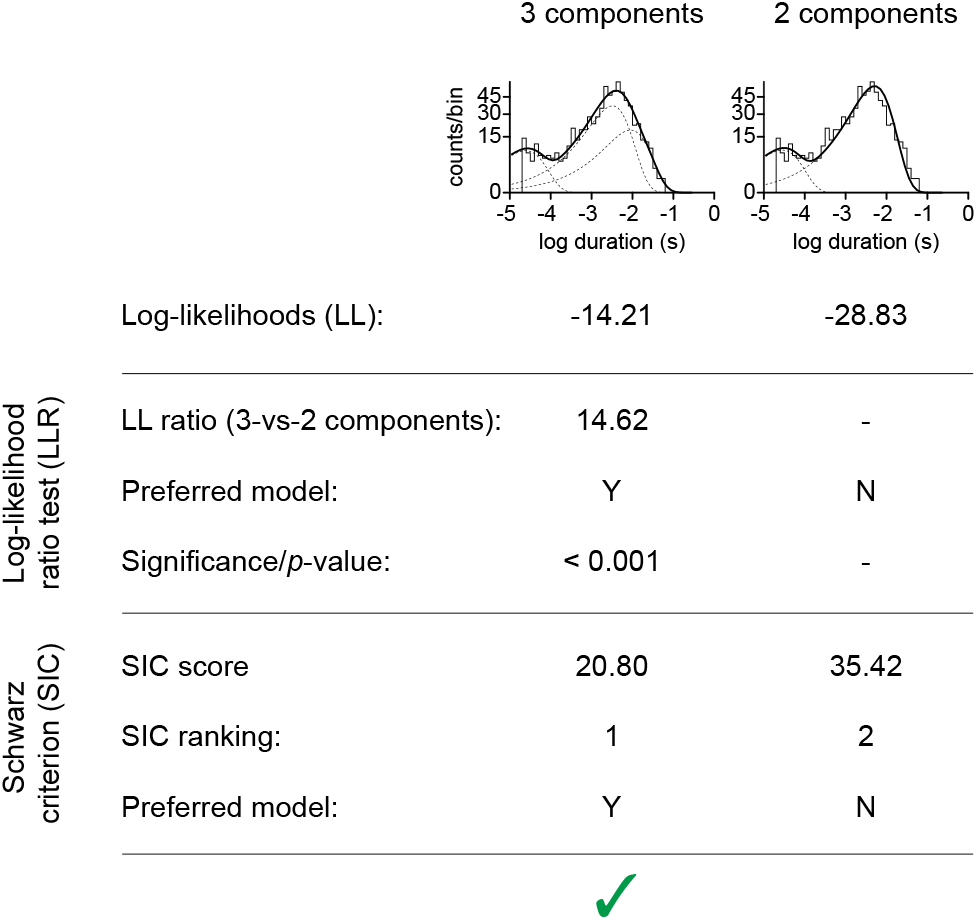
Fitting the correct number of components in the AncDGE + AncDGE-S276I mixing experiment. Open durations from cells co-transfected with AncDGE + AncDGE-S276I (alongside α and β) can be fit as the sum of two *(right)* or three *(left)* exponential components. Computed log-likelihoods for both fits are provided below each histogram. Both the log-likelihood (LL) ratio test and Schwarz information criterion (SIC) scores indicate that the three component fit is most appropriate (green checkmark). Log-likelihood ratios were calculated as LLR = LL(3 components) - LL(2 components). Signifiance was determined by comparing the LLR to the Chi-square table with 2 degrees of freedom. SIC scores were calculated as SIC = -LL + (0.5*F* ) *×* ln(*N*), where *F* is the number of free parameters, and *N* is the number of observations. Single-channel openings were elicited by 1 µM acetylcholine.

**Figure S12.**
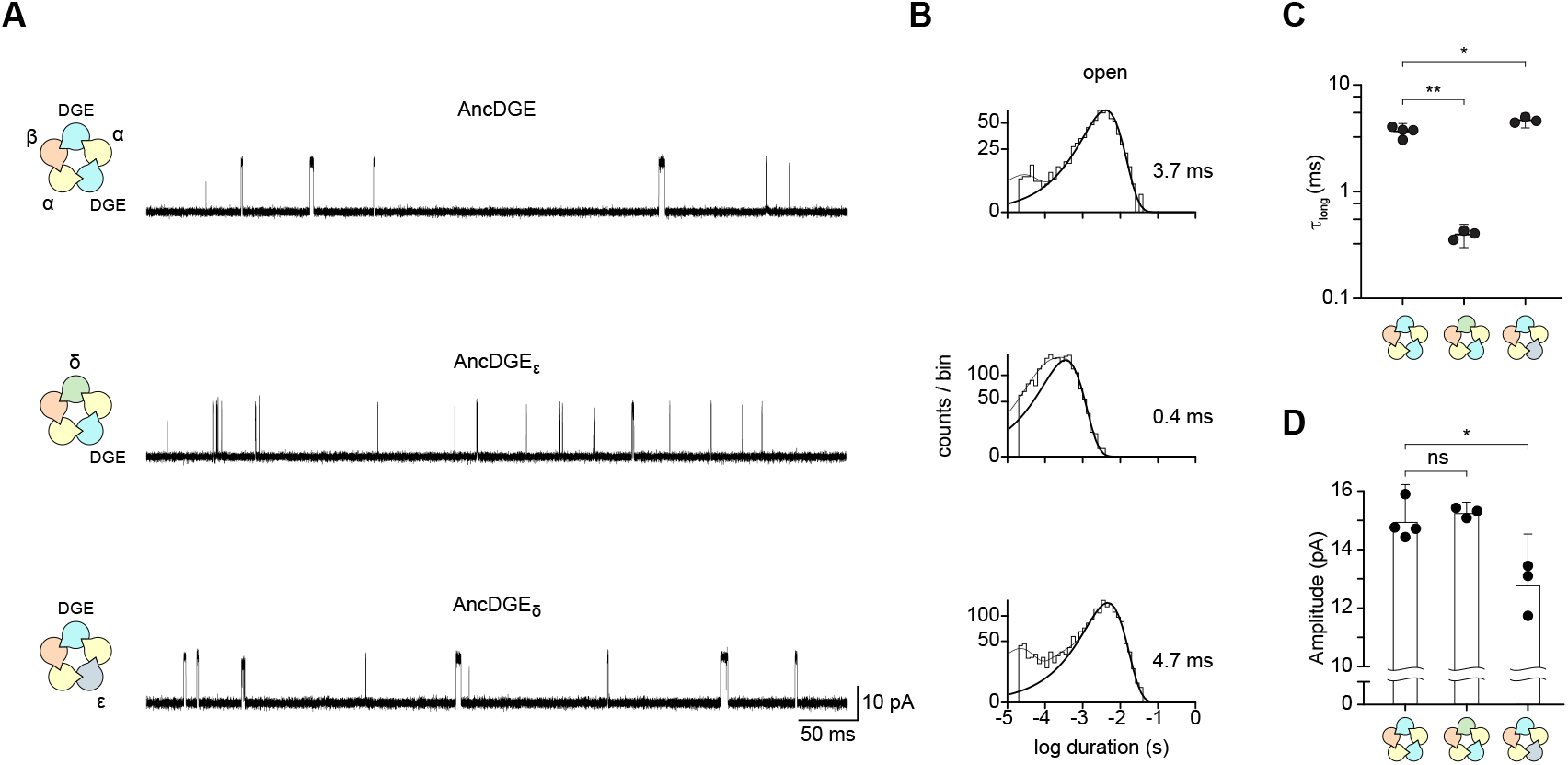
Co-transfecting α, β, and AncDGE cDNAs with an excess of δ or ε cDNA leads to AChRs incorporating only a single AncDGE subunit. **(A)** The presence of δ directs AncDGE into the ‘ε-position’ *(middle)*, while the presence of ε directs AncDGE into the ‘δ-position’ *(bottom)*. Representative single-channel traces were recorded at –120 mV in the presence of 1 µM acetylcholine. Traces are filtered at 10 kHz. **(B)** Open-duration histograms corresponding to experiments in panel A. The exponential components which represent di-liganded openings are indicated with thick black lines, with mean lifetimes (n = 3-4) shown to the right (see also Table S5). **(C)** Plot of di-liganded time constants (τ_long_). Mean ± 95 % confidence intervals are plotted with individual replicates. **(D)** Single-channel amplitudes of each AChR. A single amplitude value was obtained for each patch by averaging the amplitude of 10 individual openings. Mean ± 95 % confidence intervals are shown together with individual replicates. *p*-values for statistical differences are noted in each graph (‘^*^‘ < 0.01; ‘^**^‘ < 0.001). Subunits are colour-coded as follows: α, yellow; β, orange; δ, green; ε, blue; AncDGE, cyan.

**Figure S13.**
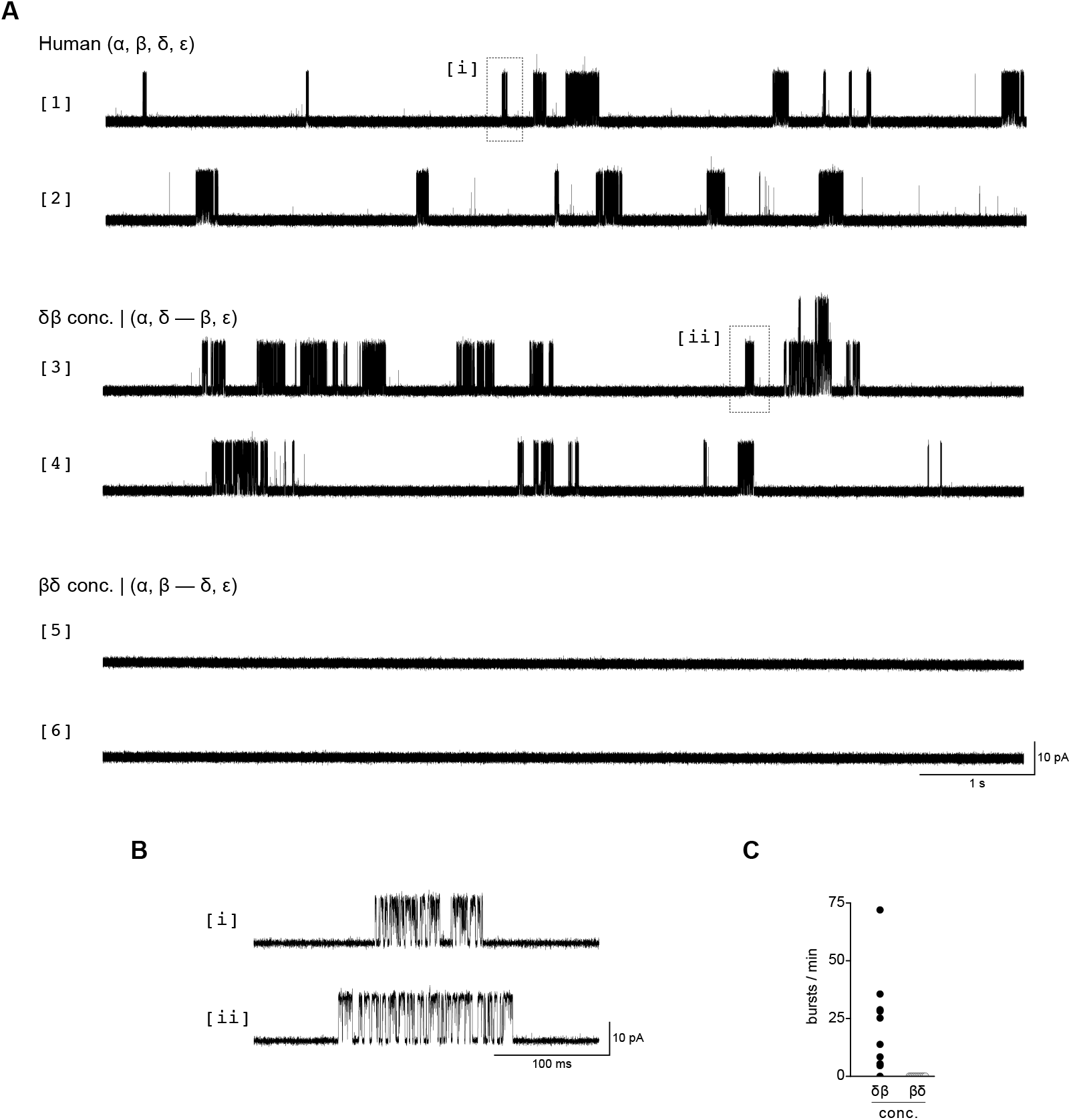
Concatenation of adult human AChRs. **(A)** Single-channel activity from cells transfected with unconcatenated AChR subunits (α, β, δ, ε; *top*; sweeps 1 and 2), α, ε, and a δβ concatemer (*middle*; sweeps 3 and 4), or α, ε, and a βδ concatemer (*bottom*; sweeps 5 and 6). Recordings were collected at 30 µM acetylcholine and –120 mV. Traces are shown filtered at 10 kHz, with openings as upward deflections. Scale bar (1 s, 10 pA) applies to all traces in panel A. **(B)** Zoom-in of boxed regions [i] and [ii] from panel A. **(C)** Single-channel activity of patches from cells expressing the δβ or βδ concatemer (alongside α and ε). The number of bursts per minute was calculated from 5 min of continuous recording (n = 10 patches).

**Figure S14.**
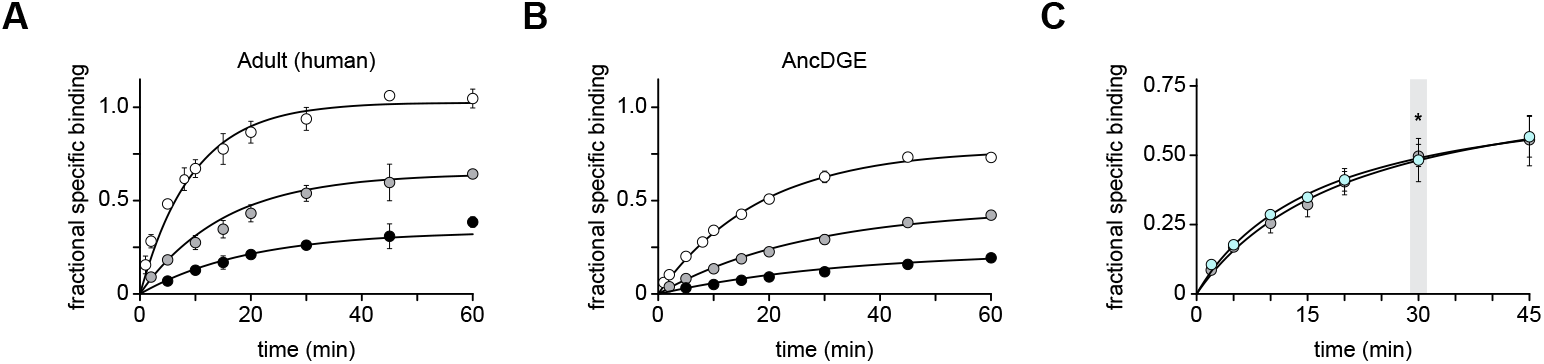
Kinetics of α-Btx binding. Cells expressing **(A)** adult human or **(B)** AncDGE-containing AChRs were incubated with increasing concentrations of [^125^I]-α-Btx (4 nM, black circles; 10 nM, grey circles; 25 nM, white circles) for the specified time. Triplicate data is shown as mean ± standard error. Association rate constants (*k*_on_) were calculated by fitting the data to the ‘Association Kinetics two ligand concentrations’ equation within GraphPad Prism (v9). *k*_on_’s were 42,307 M^-1^s^-1^ (95 % confidence interval: 36,132 M^-1^s^-1^ to 48,864 M^-1^s^-1^) and 19,553 M^-1^s^-1^ (17,615 M^-1^s^-1^ to 21,482 M^-1^s^-1^) for human and AncDGE, respectively. **(C)** 30 min incubation with 10 nM α-Btx is sufficient to occupy 50 % of available adult human binding sites (grey circles). For AncDGE-containing AChRs, the concentration of α-Btx that matches the association rate to adult human AChRs was calculated to be 21.5 nM. Accordingly, incubating AncDGE-containing AChRs (cyan circles) with 21.5 nM α-Btx yields a binding curve that is indistinguishable from that of adult human AChRs. Fractional binding is provided as mean ± standard error from three transfections. Occupancy after 30 min is marked by an asterisk (^*^), and represents the conditions under which all competition experiments were performed.

**Table S1.**
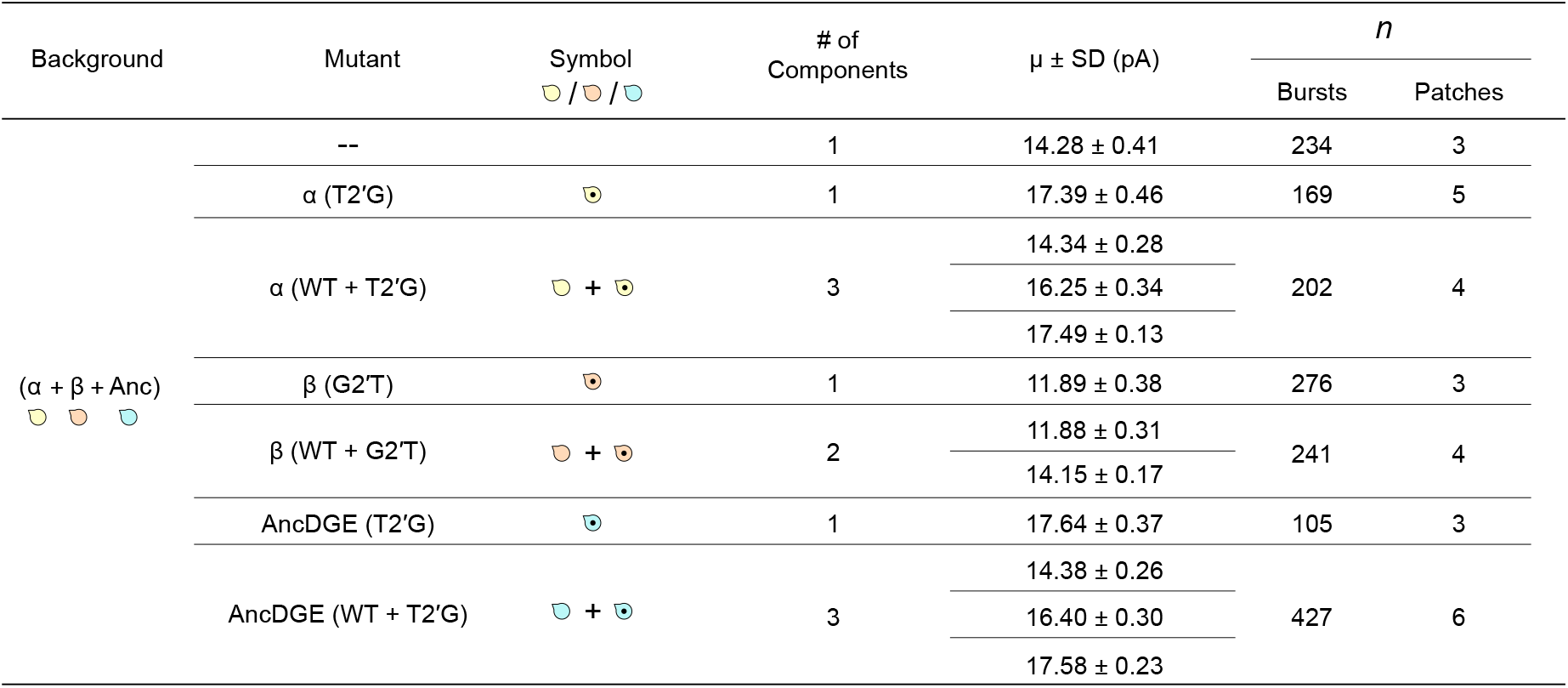
Summary of electrical fingerprinting experiments. High (HC) and low conductance (LC) subunits were transfected individually or in combination to yield channels whose amplitude depend upon the stoichiometry of the subunit in question. Mean channel amplitudes (µ) and standard deviations (SD) were extracted by fitting event-based amplitude histograms to one or more Gaussian functions. The total number of bursts in each event-based amplitude histogram, as well as the number of patches from which these bursts derive, are included. Empty subunit rosettes represent wild-type (WT) subunits without the conductance reporter mutation, while subunits with 2^*′*^ mutations are symbolized by rosettes containing a black dot.

| Background | Mutant | Symbol<br>/ / | # of Components | $\mu \pm \text{SD}$ (pA) | *n* | |
| --- | --- | --- | --- | --- | --- | --- |
|  |  |  |  |  | Bursts | Patches |
| (α + β + Anc) | -- |  | 1 | 14.28 ± 0.41 | 234 | 3 |
|  | α (T2'G) |  | 1 | 17.39 ± 0.46 | 169 | 5 |
|  | α (WT + T2'G) | + | 3 | 14.34 ± 0.28 | 202 | 4 |
|  |  |  |  | 16.25 ± 0.34 |  |  |
|  |  |  |  | 17.49 ± 0.13 |  |  |
|  | β (G2'T) |  | 1 | 11.89 ± 0.38 | 276 | 3 |
|  | β (WT + G2'T) | + | 2 | 11.88 ± 0.31 | 241 | 4 |
|  |  |  |  | 14.15 ± 0.17 |  |  |
|  | AncDGE (T2'G) |  | 1 | 17.64 ± 0.37 | 105 | 3 |
|  | AncDGE (WT + T2'G) | + | 3 | 14.38 ± 0.26 | 427 | 6 |
|  |  |  |  | 16.40 ± 0.30 |  |  |
|  |  |  |  | 17.58 ± 0.23 |  |  |

**Table S2.** Summary of competition binding experiments. Binding of acetylcholine, *d* -tubocurarine and Waglerin-1 to cells expressing either adult human or AncDGE-containing AChRs were fit to one-site and two-site competitive binding models. The best-fit model was determined using the extra-sum-of-squares F-test. IC_50_s and 95 % confidence intervals are provided for the preferred model.

| Ligand | AChR | Site 1 |  | Site 2 |  | Best-fit model | *p*-value<sup>a</sup> |
| --- | --- | --- | --- | --- | --- | --- | --- |
|  |  | IC<sub>50</sub> (μM) | 95% CI (μM) | IC<sub>50</sub> (μM) | 95% CI (μM) |  |  |
| Acetylcholine | Adult (human) | 1.2 | 0.96 - 1.6 | -- | -- | One-site | N/A<sup>b</sup> |
|  | AncDGE | 1.6 | 1.2 - 2.0 | -- | -- | One-site | 0.4985 |
| *d*-tubocurarine | Adult (human) | 0.050 | 0.0063 - 0.11 | 1.6 | 0.53 - 8.2 | Two-site | < 0.0001 |
|  | AncDGE | 0.088 | 0.075 - 0.10 | -- | -- | One-site | 0.2013 |
| Waglerin-1 | Adult (human) | 4.0 | 2.1 - 7.2 | 640 | 220 - 2500 | Two-site | < 0.0001 |
|  | AncDGE | 5800 | 3800 - 9300 | -- | -- | One-site | 0.9807 |

<sup>a</sup> *p*-values were calculated where one-site binding was set as the null hypothesis, and the two-site model being the alternate model. <sup>b</sup> No *p*-value could be calculated for acetylcholine binding to human AChRs. Two-site binding provided a poor description of the data and so the confidence interval could not be reliably computed. For this reason, single-site binding, which fit the data, was deemed the best-fit model.

**Table S3.**
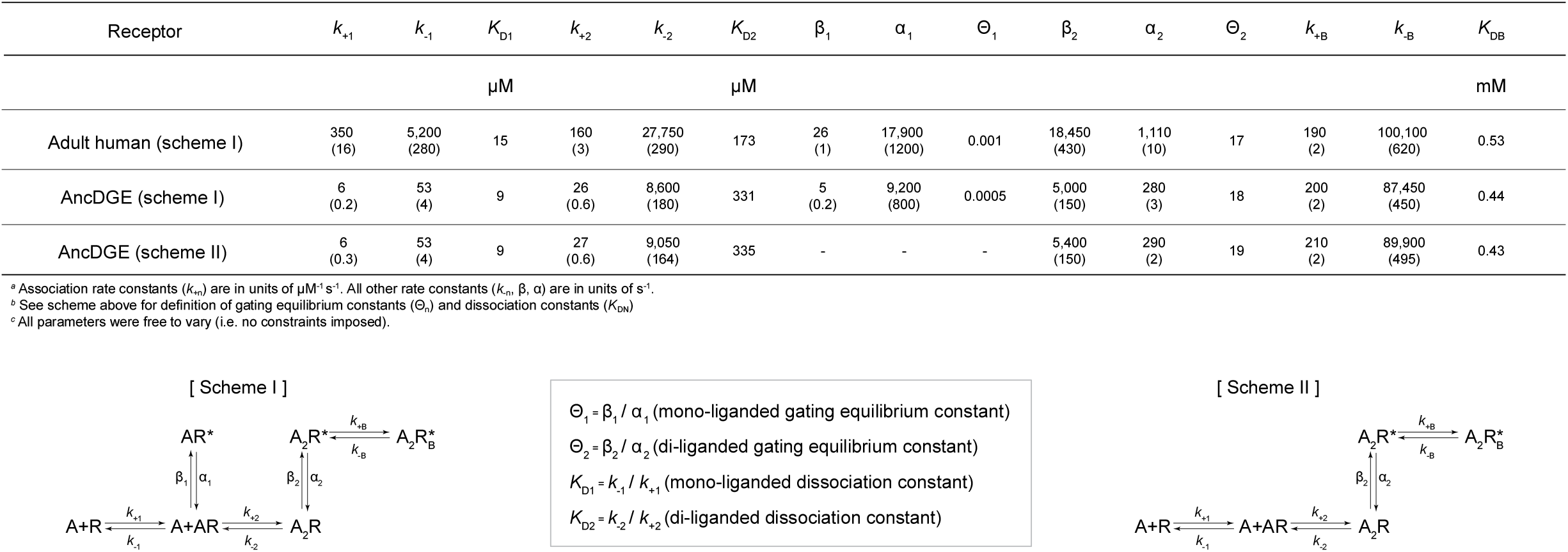
Global kinetic fitting of single-channel data. Single-channel dwells from adult human and AncDGE-containing AChRs were fit to Scheme I or Scheme II (*below* ). Rate constants and associated errors are provided (parentheses). Equilibrium constants are also defined. All data was recorded at –120 mV and digitally filtered at 10 kHz for analysis.

**Table S4.** Summary of open-channel lifetimes for adult human and AncDGE-containing AChRs at 100 nM acetylcholine. Open duration histograms were fit by a sum of three exponential components (τ_1_, τ_2_, τ_3_). Mean lifetimes (τ) ± standard error of the mean (SEM) are provided with the weighted area (± SEM) in parentheses below. All data were collected across five patches. For AncDGE, openings were pooled across these five patches due to limited activity.

| | $\tau_1$, ms<br>($a_1$) | $\tau_2$, ms<br>($a_2$) | $\tau_3$, ms<br>($a_3$) |
| --- | --- | --- | --- |
| Adult human | 0.021 ± 0.001<br>(0.21 ± 0.016) | 0.18 ± 0.020<br>(0.12 ± 0.011) | 1.40 ± 0.096<br>(0.68 ± 0.019) |
| AncDGE | 0.014<br>(0.32) | 0.080<br>(0.065) | 3.37<br>(0.61) |

**Table S5.**
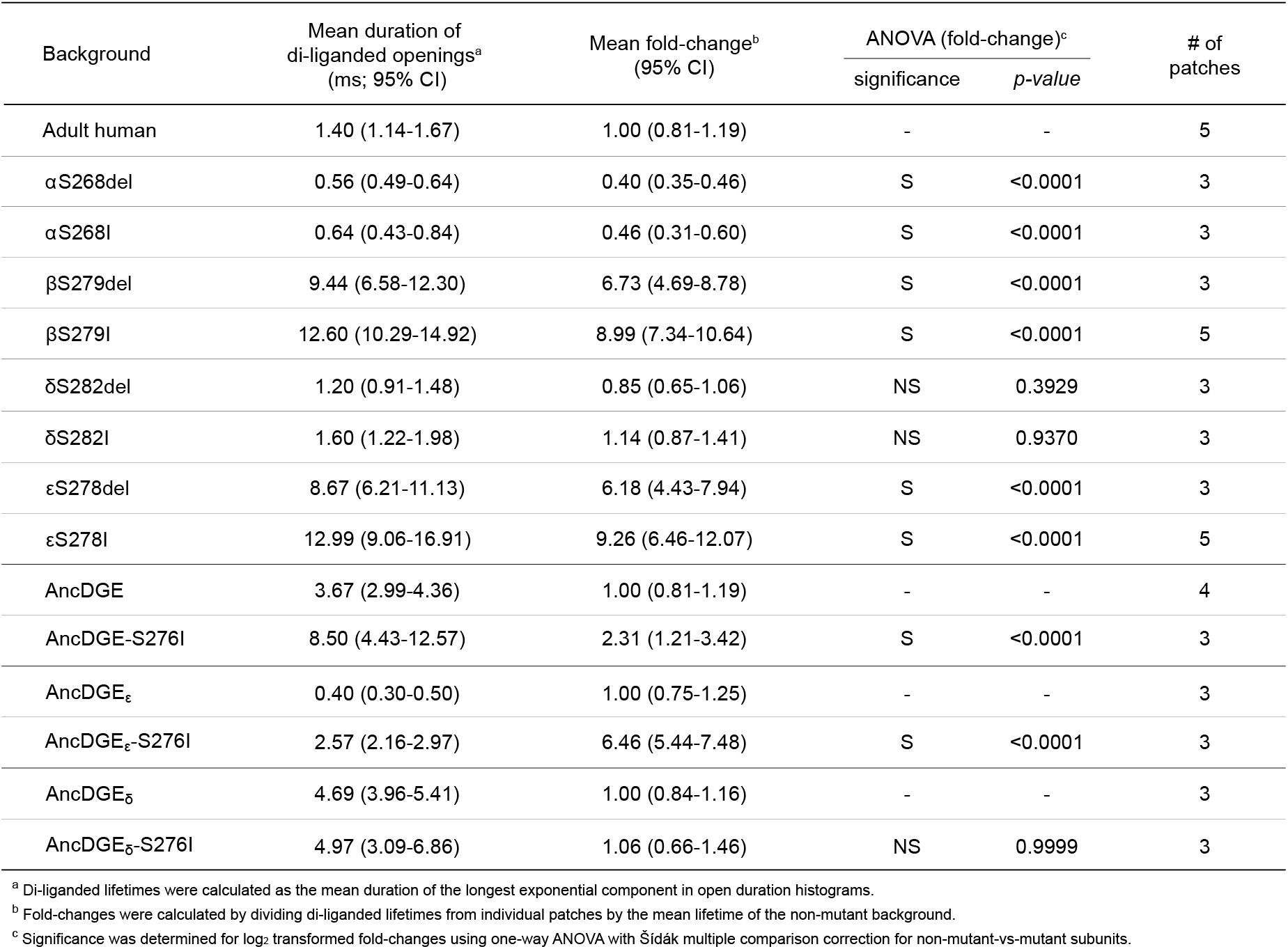
Summary statistics for human and AncDGE mutations. Shown are the mean lifetime of di-liganded openings, as well as the fold-change in this lifetime upon deletion (‘del’) or substitution of the indicated residue. Effects of mutations are relative to the unmutated backgrounds in each case (adult human and AncDGE-containing acetylcholine receptors). Also shown are the effects of the same mutation, but in the single-AncDGE subunit swap experiments (AncDGEε and AncDGE_δ_).

**Table S6.**
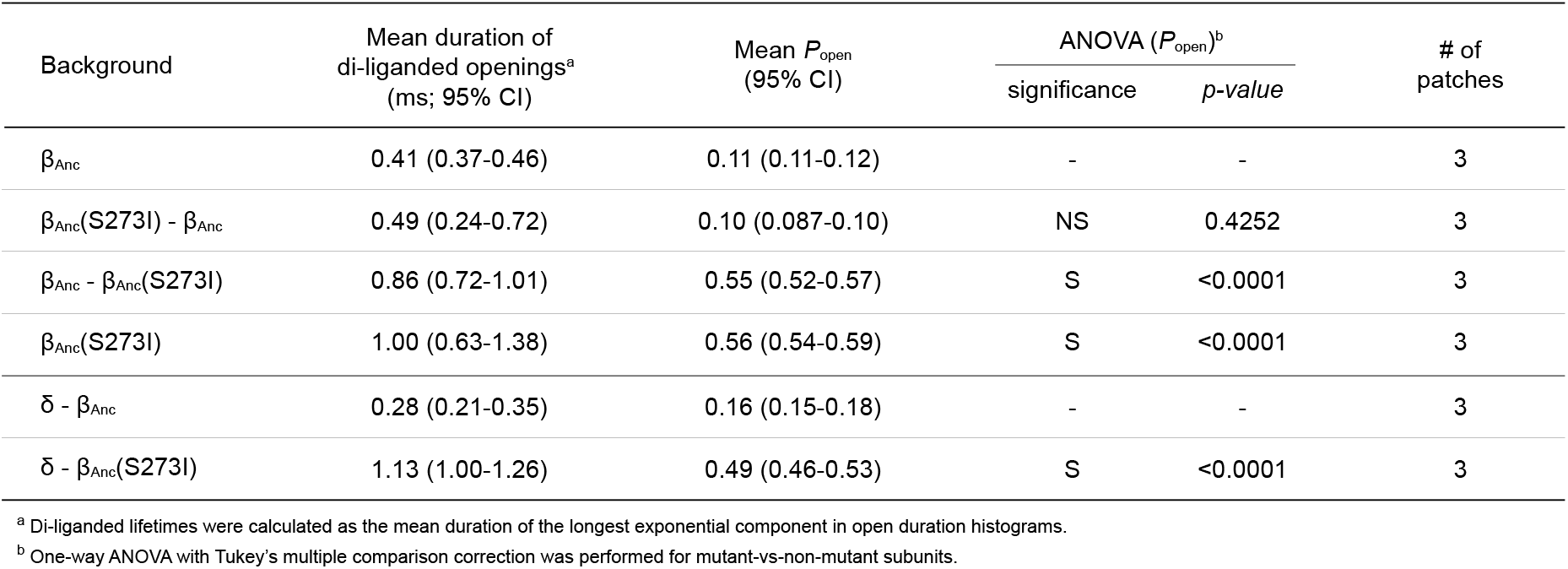
Summary statistics for β_Anc_-containing mutants. Shown are the mean lifetime of di-liganded openings, as well as the mean burst *P*_open_ upon substitution of the indicated residue. Effects of mutations are relative to the unmutated backgrounds in each case. Two subunits separated by a dash indicates a concatemeric construct.

**Table S7.**
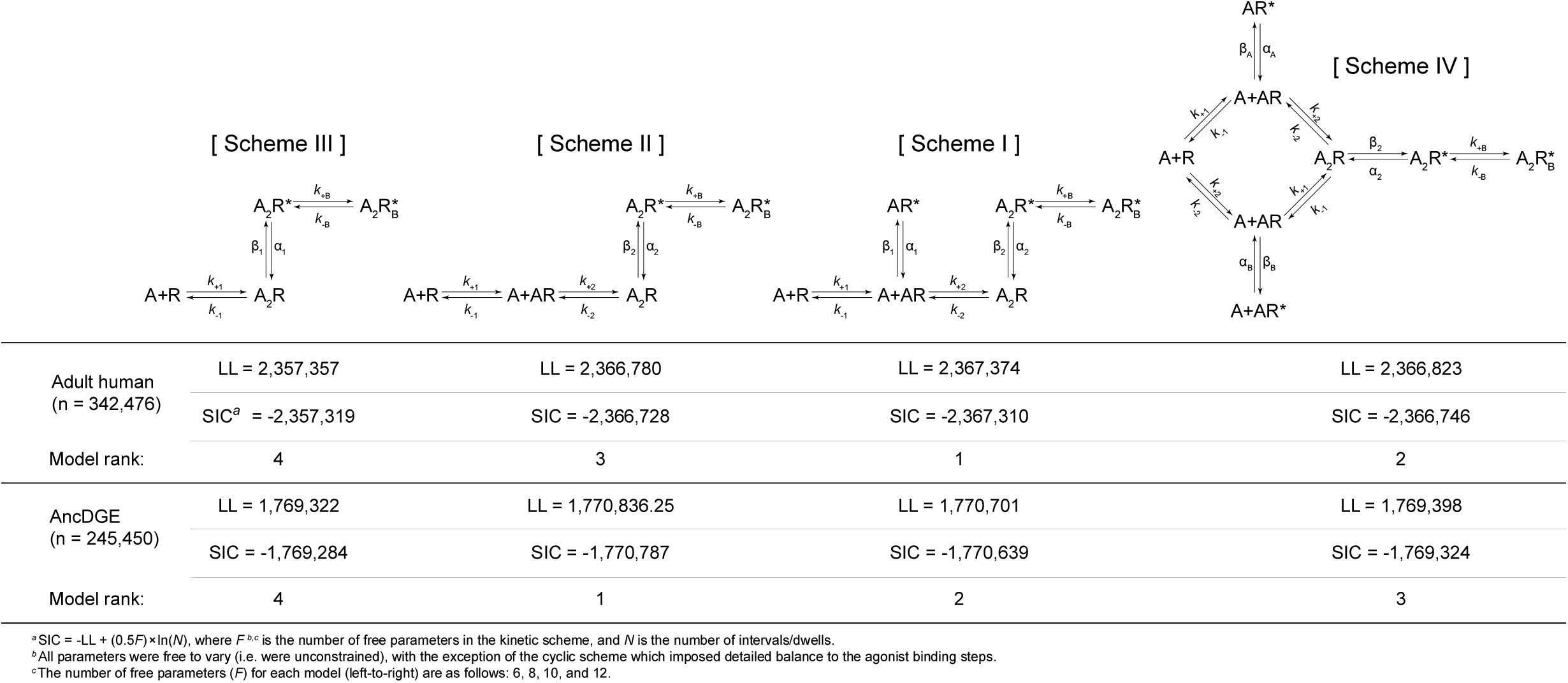
Comparison of different kinetic schemes. Open and closed dwells were pooled across several acetylcholine concentrations for adult human and AncDGE-containing AChRs, and then globally fit to the indicated schemes. Models were compared using log-likelihoods (LL) and the Schwarz information criterion (SIC). The highest-ranking model has the largest LL and smallest SIC score.

